# The role of Ca^2+^ and protein scaffolding in the formation in nature’s water oxidizing complex

**DOI:** 10.1101/2020.06.23.167841

**Authors:** Anton P. Avramov, Hong J. Hwang, Robert L. Burnap

## Abstract

The photosystem II (PSII) complex catalyzing the H_2_O-oxidation reaction of photosynthesis is highly prone to photodamage. Nature has evolved synthesis and repair mechanisms that include the photooxidative self-assembly, termed photoactivation, of the Mn_4_CaO_5_ metal cluster responsible for H_2_O-oxidation. Assembly is a multi-step light-driven process that proceeds with low quantum yield, involves a critical molecular rearrangement between light-activated steps, and is prone to photoinactivation and mis-assembly. A sensitive polarographic technique was used to track the assembly process under flash illumination as a function of the constituent Mn^2+^ and Ca^2+^ ions in genetically engineered samples to elucidate the action of Ca^2+^ and peripheral proteins. We show that the protein scaffolding that organizes this process is modulated allosterically by the assembly protein Psb27, which together with Ca^2+^, stabilizes the intermediates of photoactivation, a feature especially evident at long intervals between photoactivating flashes. Besides stabilizing intermediates, the Ca^2+^ ion is also critical to prevent photoinactivation due to inappropriate binding of Mn^2+^. Overexpression of Psb27, deletion of extrinsic protein PsbO, and excess Ca^2+^ characteristically modify these processes and retard the dark rearrangement. The results suggest the involvement of three metal binding sites, two Mn and one Ca with occupation of the Ca site by Ca^2+^ critical for the suppression of inactivation and the long-observed competition between Mn^2+^ and Ca^2+^ occurring at the second Mn site necessary for trapping the first stable assembly intermediates.

**Significance Statement:** The oxidation of water by the photosystem II is the foundation of bioproductivity on Earth and represents a blueprint for sustainable, carbon neutral technologies. Water oxidation is catalyzed by a metal cluster containing of 4 Mn and 1 Ca atoms linked via oxo bridges. The initial assembly is a complex sequential reaction harnessing the photochemical reaction center to photooxidatively incorporate Mn^2+^ and Ca^2+^ ions into the catalytic unit embedded in the protein matrix. This photoassembly is crucial for both de novo biosynthesis and as part of the ‘self-healing’ mechanism to cope with incessant photodamage that photosynthetic organisms experience. The results have implications for the natural mechanism as well as the highly desirable biomimetic devices currently envisioned for solar energy production.

## Introduction

Photosystem II (PSII) utilizes solar energy to catalyze one of the most important and most thermodynamically demanding reactions in nature: the oxidation of water into protons and molecular oxygen. The electrons extracted from the substrate water molecules are transferred through the redox-active cofactors of the photosynthetic electron transport chain eventually to reduce the electron acceptor NADP^+^, thereby forming the primary reductant for the synthesis of biomass from CO_2_ and other inorganic nutrients. Thus, the H_2_O-oxidation reaction is the basis of oxygenic photosynthetic metabolism and the primary driver of biomass accumulation on the planet (1, 2) and represents a key chemical process for the development of carbon neutral energy technologies (3, 4). Natural H_2_O-oxidation is driven by light-induced charge separation within the PSII reaction center (RC), a 700 kDa membrane protein homodimer consisting of over 20 different subunits and approximately 60 organic and inorganic cofactors [for reviews, see (1, 2)]. The catalysis of H_2_O-oxidation is mediated by a metal cluster (Mn_4_CaO_5_) buried within the protein complex at the interface between intrinsic and extrinsic polypeptides towards the luminal surface of the photosynthetic membrane (5)(**Fig. 1**). Photoexcitation of the multimeric chlorophyll (Chl) P_680_, which functions as the primary photochemical electron donor, results in the primary charge separation on a picosecond timescale into the highly oxidizing radical cation P_680_^+•^ and radical anion Pheo^-•^ (6). To minimize the backreaction or oxidation of Chl and/or neighboring proteins, the highly reactive P_680_^+•^ is re-reduced (20-250ns) by the redox active tyrosine D1-Tyr161 (Y_z_) of the D1 reaction center polypeptide located on the donor side proximal to P_680_. Meanwhile, the energized electron is stabilized by transfer from Pheo^-•^ through quinone acceptors (Q_A_ and Q_B_) and on through the remainder of the intersystem electron pathway leading to photosystem I. During the course of four consecutive charge separation events, the Mn cluster passes through a series of oxidant storage states (S-states) with the catalytic cycle balanced by removing four electrons from two bound water molecules with the release of O_2_ and four protons.

**Figure 1.**
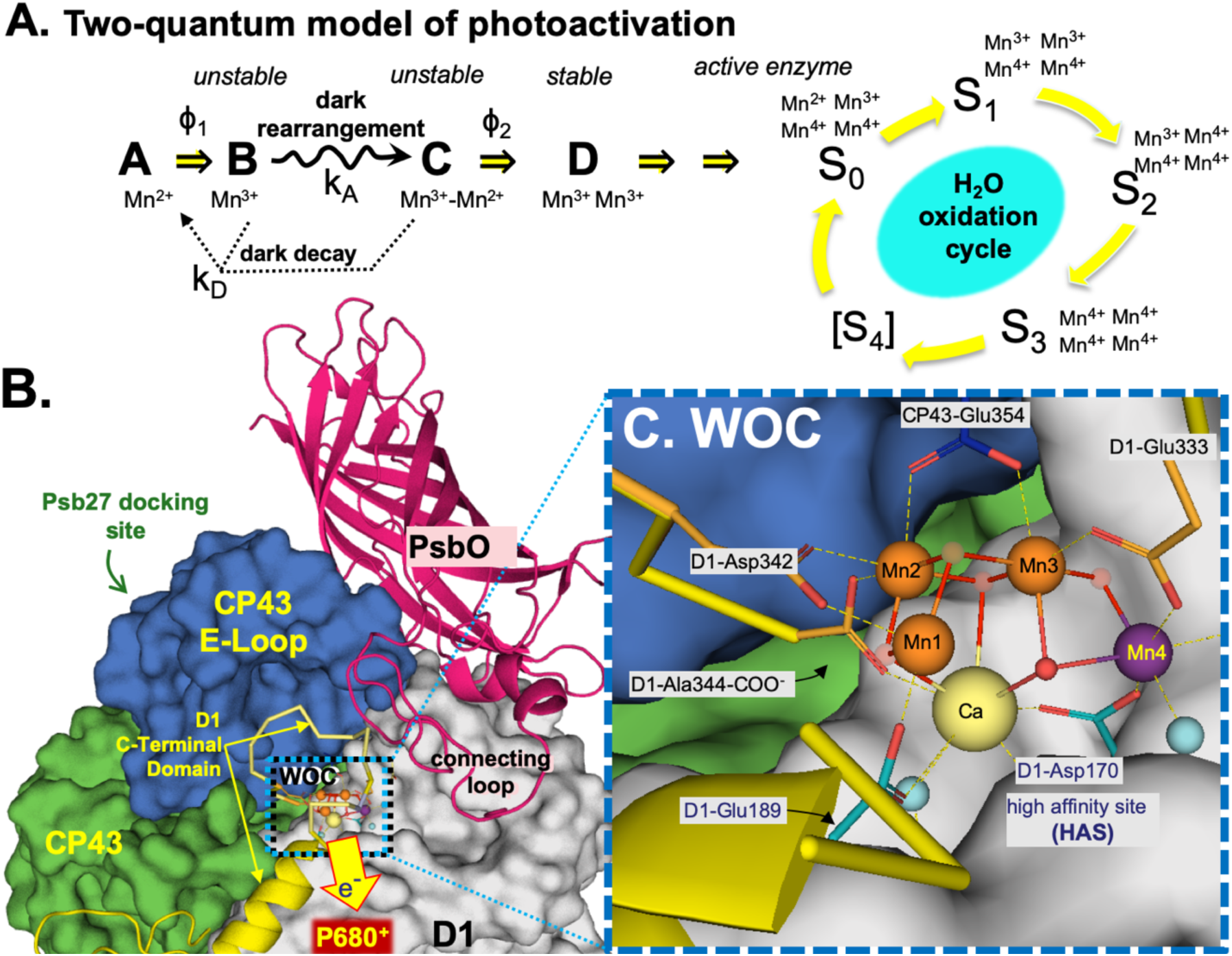
**Panel A:** Kinetic scheme of basic two-quantum mechanism of PSII photoactivation. Double arrows indicate light-activated processes with the quantum efficiencies ***ϕ***_1_ and ***ϕ***_2_ representing the first and second photooxidative events in the assembly sequence, *k*_*A*_ representing the “dark” rearrangement, and *k*_D_, representing the decay of intermediates. After the initial two Mn are photoligated, subsequent Mn appear to be added with high quantum yield. **Panel B:** The CP43 E-loop **(Blue)** interacts with the the metal binding C-terminal domain (yellow), and their coupled mobility controls access to the apo-WOC, modulated by interactions with PsbO and Psb27. **Panel C:** Catalytic Mn_4_CaO_5_ cluster at the interface between the D1 (PsbA) and CP43 (PsbC) subunits. During photoassembly, Mn^2+^ binds at the high affinity site (**HAS**) at or near the Mn4 (purple sphere) involving the D1-Asp170 carboxylate moiety (15, 16), which also ligates Ca^2+^ (yellow sphere) at the calcium site (**CAS**). Additional Mn^2+^ ions are photooxidatively incorporated via the oxo-bridges presumably derived from water ligands. A second Mn^2+^ binding site (**SMS**, see text) is critical to trap the first stable intermediate and would include amino acid ligands at one of the other three Mn locations (orange spheres).

The PSII complex is subject to incessant photodamage and a remarkable feature is its ability to undergo self-repair. Photodamage primarily occurs within PSII and much of the damage is localized in the D1 (PsbA) protein, which binds the main redox cofactors involved in photochemical charge separation. Efficient mechanisms have evolved to remove and replace damaged reaction center proteins and assemble them with their requisite cofactors (7-12). A key step in both *de novo* synthesis and the repair synthesis of PSII, is the assembly of the Mn_4_CaO_5_ core into the ligation environment of the PSII protein matrix (**Fig. 1**). Referred to as photoactivation, the assembly of the Mn_4_CaO_5_ occurs through a series of photochemical reactions that involve the oxidation of Mn^2+^ ions using the same electron extraction pathway of the fully mature PSII. Charge separation oxidizes Mn^2+^ ions to Mn^≥3+^ as the oxo-bridged multinuclear metal center forms. The multistep process begins with the binding of a single Mn^2+^ ion, (13) as its hydroxide (14) to a high affinity site (HAS) involving the D1-Asp170 carboxylate moiety (15, 16). Additional Mn^2+^ ions are photooxidatively incorporated into the growing metal cluster via the oxo-bridges that are presumably derived from water ligands of the incoming metal ions. The quantum efficiency of photosynthetic H_2_O-oxidation in the fully functional PSII is greater than 90%, whereas the photoassembly of the metal cluster is remarkably inefficient with an overall quantum efficiency below 1%, despite the fact that photooxidative assembly uses the same charge separation cofactors, notably the oxidized forms of the primary and secondary electron donors, P_680_ and Y_Z_, respectively. To account for the low quantum efficiency and for a still-unresolved light independent ‘dark-rearrangement’ that must occur between two or more light-induced charge, a so-called “two-quantum model” (**Fig. 1C**) was developed (8). Experimental support for this kinetic model is comprehensive (reviewed in (17, 18)), including, for example, direct evidence tracking the assembly process demonstrating a two-quantum requirement, and that part of the overall inefficiency is due to the competition between the slow and/or inefficient assembly steps and back-reactions of the charge separated state (19). Nevertheless, evidence regarding the structure and chemistry of the intermediates and the nature of the dark rearrangement has remained scarce.

Calcium is a critical cofactor in the process of H_2_O-oxidation by PSII because it mediates the delivery of substrate water to a Mn coordination site for oxidation and dioxygen formation. While early studies suggested that Ca^2+^ is not required for photoassembly, there is now general agreement that Ca^2+^ is vital (20-22). Nevertheless, the mechanism of how Ca^2+^ facilitates the photoassembly remains obscure. Binding of Mn^2+^ to the HAS with simultaneous binding of Ca^2+^ to an adjacent binding site facilitates the formation of the [Mn^2+^-(OH)-Ca^2+^] complex by inducing deprotonation of a water ligand of Mn^2+^. Calcium lowers the pK_a_ for water ligand, which is controlled by a nearby base B^-^ that serves as a primary proton acceptor with a p*K*_a_ dependent on Ca^2+^ bound to its effector site (23). The deprotonation of the intermediates and the tuning of pK_a_s facilitated by Cl^-^ binding is critical for assembly (24). The absence of Ca^2+^ during photoactivation leads to the formation of non-catalytic, multinuclear high valent Mn species (“inappropriately bound Mn”) that inactivates the PSII complex (22). At the same time, high concentrations of the Ca^2+^ cofactor diminishes the efficiency of photoactivation. These results indicate that Ca^2+^ and Mn^2+^ compete for each other’s binding sites, leading to an optimality relationship in their relative concentrations during photoactivation (10-12).

Three extrinsic polypeptides, PsbO, PsbV, and PsbU, serve to enclose and stabilize the cyanobacterial water oxidation complex (WOC), whereas plants and certain eukaryotic algae possess PsbO, PsbQ, and PsbP, [for review see, (25)]. The extrinsic polypeptides prevent the reduction of the Mn cluster by exogenous reductants (26-28) and help retain the Ca^2+^ cofactor, which is otherwise prone to loss during the catalytic cycle (29). However, by enclosing the WOC, accessibility of ions for the photoactivation of the Mn_4_CaO_5_ may be restricted. Indeed, the most efficient *in vitro* photoactivation procedures either explicitly or coincidentally involved the biochemical removal of the extrinsic proteins and genetic deletion of the most evolutionarily conserved extrinsic protein, PsbO, increases the quantum efficiency of photoactivation (17). Proteomic analysis of highly intact cyanobacterial PSII revealed novel proteins, including a small protein designated Psb27, apparently associated with PSII sub-populations (30). Psb27 was identified as a lipoprotein associated with inactive PSII monomers that prevents binding of PSII extrinsic subunits (PsbO, PsbU and PsbV) to the premature PSII (31), keeping the active site “open” and thereby maintaining a sufficient diffusion rate of Mn^2+^, Ca^2+^ and Cl^-^ ions thus acting as a molecular chaperone for successful photoactivation (32) and does so through a specific interaction with the E-loop of CP43, which is a lumenal domain of PSII that directly interacts with the Mn_4_CaO_5_. Overall, Psb27 appears to be strictly associated with organisms possessing PSII and is important for its assembly, (33-36) yet its precise role in facilitating assembly remains to be resolved. The application of *in vitro* photoactivation procedures using genetically tractable cyanobacteria has not been described and here we describe such a system and use it for the analysis of the role Ca^2+^ and Psb27 in the assembly of the Mn_4_CaO_5_.

## Results

### Mn^2+^ and Ca^2+^ competition during photoassembly leads to decreased quantum efficiency and yield during photoactivation of Synechocystis membranes

We developed a procedure for isolating Mn-depleted thylakoid membranes from *Synechocystis* sp. PCC6803 (hereafter, *Synechocystis*) that permits *in vitro* studies with control of the photoactivation conditions, such as pH and ion composition, in an experimental system that also allows facile genetic modification. Importantly, the membranes retain the native electron acceptor system so that artificial electron acceptors are not necessary as in previous *in vitro* experimental systems. Extracted membranes showed substantial restoration of photosynthetic activity (40%) (**Fig. S3A, Table S1**) consistent with published yields in plant preparations (see (12) for quantitative analysis), and most importantly, display similar kinetic features of photoactivation compared to plant membrane preparations and to *in vivo* photoactivation experiments in *Synechocystis* (**Fig. S4**). We focused on the role of Ca^2+^ in photoactivation and address the question as to how the extrinsic proteins modulate the demand for both Mn^2+^ and Ca^2+^. As shown previously, photoactivation requires presence of both Mn^2+^ and Ca^2+^ cations for the assembly of PSII and post-addition of Ca^2+^ only resulted in a very small increased yield of photoactivation in the dark (**Fig. S3B)**. Post-addition of Sr^2+^ or Mg^2+^ cations in the dark, did not result in a significant yield of photoactivation. Thus, our results concur with the general conclusion that Ca^2+^ is required for the assembly process (20-22).

In plant preparations, optimal photoactivation *in vitro* (11, 12, 37) and *in organello* (10) requires an optimal Ca^2+^/Mn^2+^ ratio, with an excess of Ca^2+^ relative to added Mn^2+^, which reflects a competition between the ions for their respective binding sites (12). To establish whether similar kinetic features operate in cyanobacteria, the Ca^2+^ concentration dependence of photoactivation was performed at two fixed Mn^2+^ concentrations, 250µM or 500µM (**Fig. 2A & B**). These concentrations of Mn^2+^ should saturate its binding to the HAS, (11) but should still allow observation of the predicted competition between the two cations observed in plant preparations (10-12). A small level of photoactivation can be seen in the absence of added Ca^2+^ ions, which could be explained by trace amounts of residual Mn^2+^ and Ca^2+^ ions remaining in the extracted thylakoid preparation, since the level of this activity could be minimized but never completely eliminated by extensive washing of membranes using Chelex-treated buffer. Interestingly, the yield of photoactivation in the absence of Ca^2+^ only, while low, is nevertheless higher than in the absence of Mn^2+^ only (**Fig. S5**), which probably reflects a sub-optimal Ca^2+^/Mn^2+^ ratio leading to photoinactivation (*vide infra*). For all the samples containing 250µM Mn^2+^ in photoactivation buffer, the maximum yield of photoactivation was observed at 600-700 flashes with the half-maxima at occurring with approximately 150-200 flashes (**Fig. 2A**). In terms of maximal yield and quantum efficiency of photoactivation, the optimum Ca^2+^ concentration for photoactivation at 250µM Mn^2+^ occurs at 10mM corresponding to Ca^2+^/Mn^2+^ ratio of 40:1. The decrease in O_2_ evolution at lower Ca^2+^ concentrations is likely caused by the competitive binding of Mn^2+^ to the Ca^2+^ binding site. In the context of earlier findings, this would prevent the formation a bridged species [Mn^3+^-(OH)-Ca^2+^] that is proposed to be crucial intermediate during the assembly, (23) and perhaps leading to the formation of ‘inappropriately’ bound high valence, multinuclear Mn deposits (22). To better understand the role of ion competition and to test the hypothesis that Mn^2+^ binding at the Ca^2+^ site leads to photoinactivation of PSII during photoassembly, we carried out the same experiment, but doubling Mn^2+^ concentration to 500µM (**Fig. 2B**). For the optimum photoactivation at 500µM of Mn^2+^ the requirement for Ca^2+^ increased two-fold. Interestingly, sample containing 20mM and 40mM of Ca^2+^ in photoactivation buffer started to show signs of photoinactivation observed earlier at low Ca^2+^ concentration at 250µM Mn^2+^. From these results we can conclude that an excess of Mn^2+^ ions leads to the decreased yield of PSII photoactivation due to photoinactivation, while excess of Ca^2+^ ions does not have such effect.

**Figure 2.**
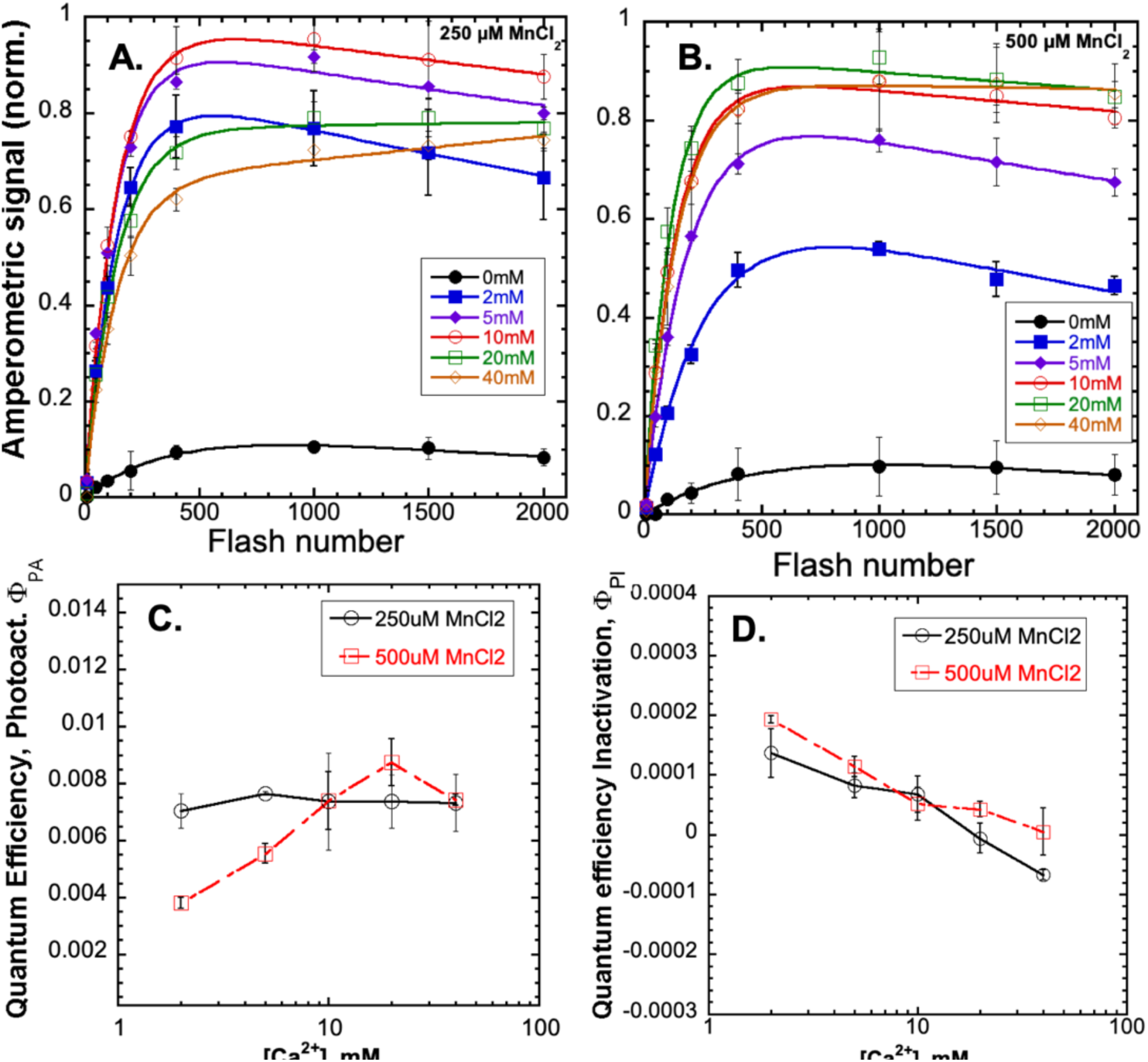
Calcium dependence of photoactivation under sequence of single turnover flashes of HA-extracted thylakoid membranes from WT control at 0mM (black closed circle), 2mM (blue closed square), 5mM (purple closed diamond), 10mM (red open circle), 20mM (green open square), and 40mM (orange open **diamond**) of CaCl_2_ combined with 250 μM (**Panel A**) or 500 μM MnCl_2_ (**Panel B**). **Panel C:** Overall quantum efficiency of photoactivation (*Φ*_*PA*_**) Panel D:** quantum efficiency of inactivation (*Φ*_*PI*_). Data were fit to equation 1 for parameter estimation (see text for details). Error bars represent standard deviation n ≧3.

To analyze the role of Ca^2+^ concentration on quantum efficiency of photoactivation and photoinactivation as a function of flash number and to better understand inhibitory effect of low Ca^2+^ to Mn^2+^ ratio, data described in **Fig. 2** was fit to a double exponential equation:

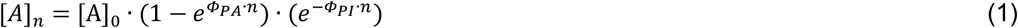

The equation accounts for the a progressively smaller pool of apo-centers during the flash sequence progression as more centers become photoactivated (9) combined with a term that represents the loss of centers due to photoinactivation processes (12). Here, [A]_*n*_ represents the yield of active centers on the *n*th flash, whereas [A]_0_, is the concentration of apo-PSII centers prior to the photoactivation, *Φ*_*PA*_, represents the overall efficiency of the multiphoton assembly process. The photoinactivation term, *Φ*_*PI*_, represents the quantum efficiency of irreversible photodamage or, alternatively, the formation of inactive centers due to ‘inappropriately bound’ Mn as a consequence of supra-optimal Mn^2+^ concentrations. This is also seen with varying [Mn^2+^] at fixed [Ca^2+^], where there is a strong dependence of *Φ*_*PI*_ on [Mn^2+^] (**Fig. S6**). The corresponding estimates of parameters are plotted as a function of the Ca^2+^/Mn^2+^ ratio (**Fig. 2C & D**). While the yields of photoactivation increase with [Ca^2+^] up to 10mM and 20mM at 250µM Mn^2+^ and 500µM Mn^2+^, respectively, higher [Ca^2+^] decreased yields **(Fig. 2A & B)**. Interestingly, at 250µM Mn^2+^ the quantum efficiency of photoactivation, *Φ*_*PA*_, is relatively unaffected by Ca^2+^ concentration throughout the range tested (**Fig. 2C**). In contrast, at the higher (500µM Mn^2+^) concentrations, *Φ*_*PA*_ is strongly dependent on Ca^2+^ availability, perhaps reflecting competitive occupation of Mn^2+^ in the Ca^2+^ effector site. As shown in **Fig. 2C**, at low [Ca^2+^], the *Φ*_*PA*_ is low, reaches a maximum at 20mM coinciding with overall apparent optimum (**see Fig. 2B**), and declines at higher [Ca^2+^]. Overall, higher abundance of Mn^2+^ ions inhibits the assembly through competition with Ca^2+^ (10-12). However, the difference between the unaffected *Φ*_*PA*_ at 250µM Mn^2+^ versus its being affected at 500µM in **Fig. 2C** is intriguing and suggests a complex interplay of binding constants. The HAS likely remains occupied by Mn^2+^ (K_D_ < 10µM), (11, 13, 15) with both the 250 and 500µM Mn^2+^ experiments, even with comparatively high [Ca^2+^]. However, at the higher [Mn^2+^] the results indicate that the Ca^2+^ effector site is substantially occupied by Mn^2+^ leading to inactivation, consistent with previous kinetic analysis (12). It is unclear at which step(s) the replacing of Ca^2+^ ion with Mn ion is inhibitory, however we find that Ca^2+^ can even be replaced with Mn in assembled PSII, producing a light-dependent inactivation during the catalytic turnover of the S-state cycle of the intact Mn cluster (**Fig. S7**). This fits with the observation that Ca^2+^ is more readily released in the higher S-state, presumably due to charge repulsion (29, 38). Accordingly, this replacement by Mn^2+^ potentially results *not only* in a failure to advance in photoassembly, but could lead to a greater frequency of inactivation mitigated by high [Ca^2+^] evidenced by the declining *Φ*_*PI*_ with increasing [Ca^2+^] (**Fig. 2D)**. The optimal Ca^2+^/Mn^2+^ ratio for overall photoactivation thus reflects a balance between binding of Ca^2+^ at its effector site, which prevents photoinactivation due to inappropriate binding of Mn^2+^, (22) but not so high as to out-compete Mn^2+^ at a Mn site preventing photooxidative Mn incorporation (11-13). But which Mn site? Based upon the fact that *Φ*_*PA*_ at lower [Mn^2+^] is independent of [Ca^2+^] in the range tested, as well as previous estimates of Mn^2+^ affinity at the HAS, we conclude that the yield-limiting competition between Ca^2+^ at a Mn^2+^ site occurs not at the HAS, but rather at a second Mn site (SMS) involved in the photoactivation pathway. This is consistent with the results of the effect of [Ca^2+^] on the rate of the dark rearrangement step, k_A_ (next section).

### Calcium stabilizes intermediates of photoactivation and prolongs the dark rearrangement time

To gain further insight on the role of Ca^2+^ in photoactivation, experiments were performed varying the interval between photoactivating light flashes for a fixed number of flashes to test the hypothesis that Ca^2+^ influences the stability of the intermediates of the photoassembly process **(Fig. 3**). According to the two-quantum model of photoactivation (**Fig. 1C**) the initial photooxidation of Mn^2+^ (state **“A”**) produces the first unstable intermediate **“B”**, followed by a light-independent ‘rearrangement’ step leading to the formation of a second unstable intermediate **“C”**, which is a configuration capable of productively utilizing the second light quantum to form the first stable intermediate, **“D”** (8). Short intervals produce low yields since not enough time has elapsed for completion of the dark rearrangement. Long intervals between the charge separations causes the decay of intermediates resulting low yields of active centers (7, 8). This results in a bell-shaped curve recording the yield of photoactivation due to a fixed number of flashes given at different flash intervals from 50ms to 10000ms (**Fig. 3**). Estimations of the parameters under each condition **(Table 1)** describing the dark rearrangement, *k*_A_, and the decay of intermediates, *k*_D_, were determined by deriving kinetic parameters from the rising and falling slopes of the bell-shaped curve in plots of photoactivation as a function of the flash interval curve fitted to the equation (2):

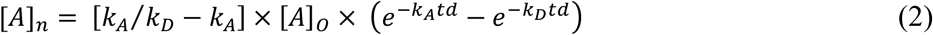

**Table 1.** Dark rearrangement, k_A_ and decay of intermediates, k_D_, parameters^a^ characteristic of photoactivation of membranes from WT, ΔpsbO, and 27OE strains.

| Strain / [Ca<sup>2+</sup>] | $k_A(s^{-1})$ ($t_{1/2}$) | $k_D(s^{-1})$ ($t_{1/2}$) | fit quality, r |
| --- | --- | --- | --- |
| **WT control 2mM Ca<sup>2+</sup>** | 2.4 (288ms) | 0.134 (5.17s) | 0.99 |
| **WT control 10mM Ca<sup>2+</sup>** | 2.8 (244ms) | 0.095 (7.29s) | 0.98 |
| **WT control 40mM Ca<sup>2+</sup>** | 1.6 (433ms) | 0.051 (13.5s) | 0.99 |
| **$\Delta psbO$ 2mM Ca<sup>2+</sup>** | 1.9 (364ms) | 0.104 (6.66s) | 0.99 |
| **$\Delta psbO$ 10mM Ca<sup>2+</sup>** | 2.2 (315ms) | 0.072 (9.62s) | 0.99 |
| **$\Delta psbO$ 40mM Ca<sup>2+</sup>** | 1.5 (462ms) | 0.049 (14.1s) | 0.99 |
| **27OE 2mM Ca<sup>2+</sup>** | 1.5 (462ms) | 0.130 (5.33s) | 0.98 |
| **27OE 10mM Ca<sup>2+</sup>** | 1.4 (495ms) | 0.085 (8.15s) | 0.99 |
| **27OE 40mM Ca<sup>2+</sup>** | 1.8 (385ms) | 0.027 (25.6s) | 0.99 |

<sup>a</sup> Data were fit to equation 2 (see main text) to estimate the dark rearrangement constant, $k_A$ , and the decay of photoactivation intermediates “B and C”, $k_D$ (see Fig. 1 for model, and Table 1 for estimated values for $k_A$ and $k_D$ ). Error bars represent standard deviation $n \geq 3$ .

**Figure 3.**
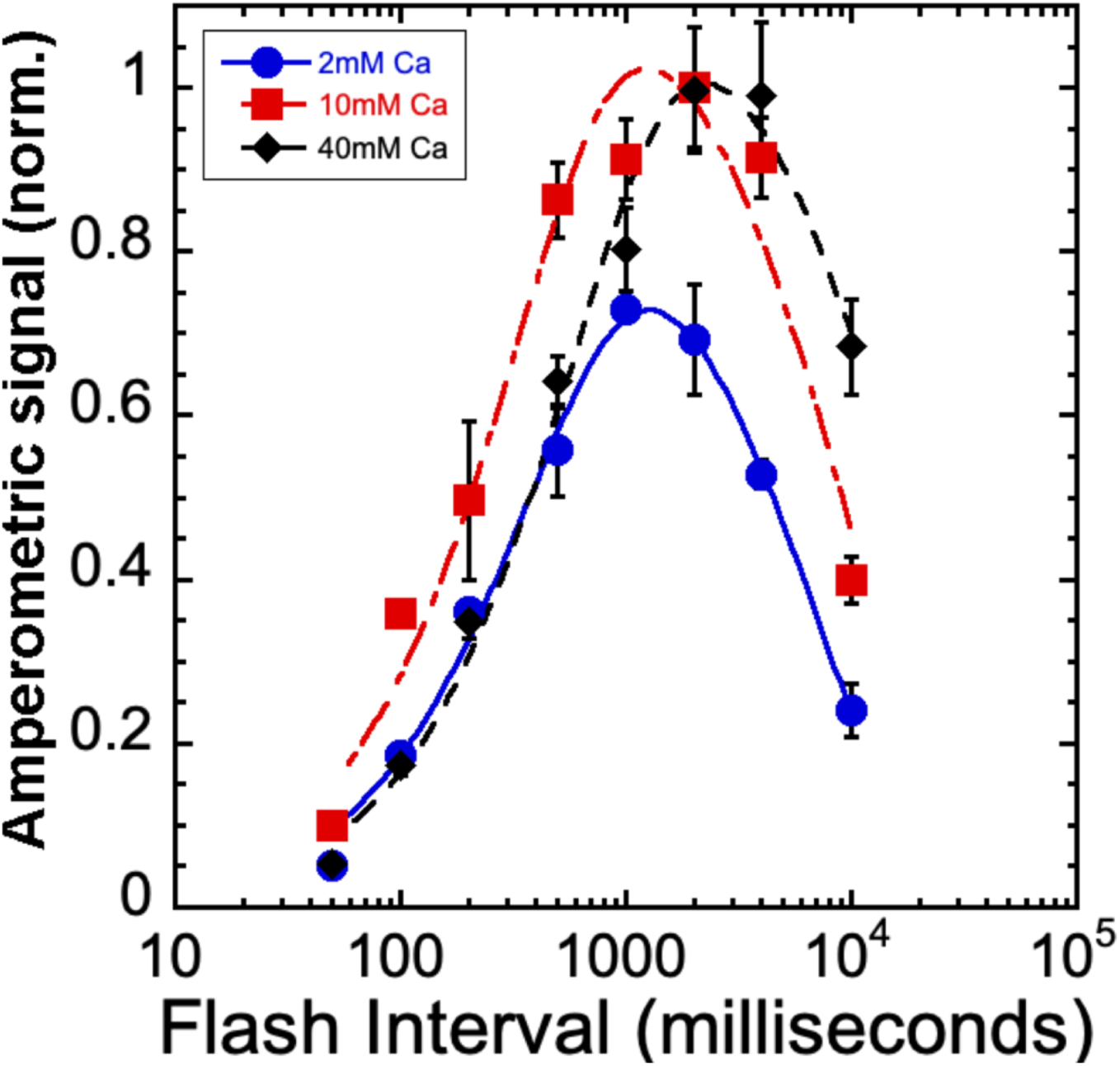
Photoactivation yields as a function of flash interval at different calcium concentrations. Sequences of 150 Xe flashes applied at different flash intervals to HA-extracted thylakoid membranes from WT control and oxygen yields measured on a bare platinum electrode. Plots correspond to samples containing 2mM (blue circles), 10mM (red squares), and 40mM (black diamond) of CaCl_2_ a fixed [Mn^2+^] = 250 μM. Error bars represent SD with n ≧3. Data was fit to equation 2 (see main text) to estimate the dark rearrangement constant, k_A_, and the decay of photoactivation intermediates “B and C”, k_D_ (see Fig. 1 for kinetic model and **Table 1** for estimated values for k_A_ and k_D_).

The overall shape of the resultant bell-shaped curves closely resembles previous results using higher plant material with estimated values for dark rearrangement (k_A_, **B→C**) and the decay of intermediates (k_D_, **B** & **C**) are in the range of 200-400 ms and 7-14 seconds, respectively in our experiments (**Table 1**). These values are slower than with spinach PSII preparations, but in the same proportions with the dark rearrangement, k_A_, typically 10-fold faster than the rate of decay, k_D_, (reviewed in (18)). These differences may be attributed to the different source of experimental material and/or choice of buffers (e.g. the sucrose concentration is much higher than typically used). Depending upon the [Ca^2+^], different optima for maximal yields of photoactivation were observed with increasing Ca^2+^ giving increasingly longer optima for the length of the dark period between flashes (**Fig. 3**). Photoactivation with 40mM Ca^2+^ enhanced yields at the longest interval tested, (10 sec) again suggesting that Ca^2+^ stabilizes the intermediates of photoactivation. Accordingly, the fit values for the decay of intermediates *k*_D_, are shifted to slower rates of decay (**Table 1**). The stabilizing effect of Ca^2+^ on photoactivation intermediates contrasts with the existence of the optimal [Ca^2+^]/[Mn^2+^] ratio for the net yield under repetitive flashing at a constant interval (500 ms). The second parameter estimated by the flash interval experiment is the dark rearrangement, *k*_A,_ (**B→C, Fig. 1**). This is the time needed between flashes before subsequent flashes become productive, which is experimentally reflected by diminished yields of photoactivation at short flash intervals (**Fig. 3**). Remarkably, the rearrangement appears to take longer at high [Ca^2+^] concentrations **(Table 1)**.

Our analyses of the preceding results leads us to conclude that at high [Ca^2+^], assembly occurs, albeit sub-optimally, due to competition between metals for their respective sites as shown before, (10-12), yet high [Ca^2+^] also allows assembly at long flash intervals due to stabilization of intermediates **“B”** and/or **“C” (Fig. 1)**. On the other hand, at low [Ca^2+^], photoactivation yields are also decreased **(Fig. 3)** due to photoinactivation. The protective function of Ca^2+^ is important also in PSII centers that already have been assembled **(Fig. S7)** consistent with the proposed ‘gate-keeper’ function observed with intact PSII preparations (28, 39). Thus, we conclude that Ca^2+^ is important for assembly because it stabilizes intermediate **“B”** and/or **“C”** and that it prevents inactivation and failure to advance due to ‘inappropriate’ binding of Mn^2+^ into the Ca^2+^ site during assembly (12, 22). Besides its “gate-keeping” role, Ca^2+^ also can block necessary binding of the second Mn^2+^ especially at very high [Ca^2+^]/[Mn^2+^] ratios. This accounts for the slowing of the very slow dark rearrangement (**B→C).** While sufficient concentrations of Ca^2+^ increase the chance of successful formation of intermediate **“C”**, an excess of Ca^2+^ competes with the binding of the second Mn^2+^, thereby delaying the time before the second Mn^2+^ ion can occupy its site for photooxidation. This competitive inhibition extends the time before the rearranged state can be trapped (**C⇒D**) until the competing Ca^2+^ is replaced with Mn^2+^ enabling the photooxidative formation of stable intermediate **“D”**. This, in turn, suggests that the dark rearrangement consists of a molecular reorganization (e.g. conformational change) that is only fruitful if a second Mn^2+^ bound at its correct site.

### Role of PsbO and Psb27 during photoactivation

The PSII assembly cofactor protein, Psb27, which appears to facilitate diffusional access to the WOC assembly site (31, 32, 36), does so by interacting with the E-loop of CP43 to allosterically modify its conformation (35), which is hinged (18) in a way that opens the WOC for greater diffusion into the sites of cofactor binding (32). Since the natural steady-state abundance of Psb27 in the cell likely evolved to cope with assembly and repair of only a fraction of centers from the total population of PSII in the cell, then there may not be enough copies of Psb27 to stoichiometrically service all PSII centers upon quantitative removal of the Mn cluster experimentally produced by HA-extraction. Similarly, diffusion of Mn^2+^ and Ca^2+^ ion could also be limited by the presence of the extrinsic proteins, impeding photoassembly as previously suggested from *in vivo* experiments with mutants (32, 40). To address these considerations, the photoactivation of two *Synechocystis* mutants, Δ*psbO* lacking PsbO, (41) and a strain overexpressing Psb27 (*27OE*) **(Fig. S8)**, were investigated. Importantly, the more open configurations of the WOC in both strains enabled a net recovery of O_2_ evolution under continuous illumination that was more than 50% higher than in the WT-control (**Table S2**). Additionally, the optimal [Ca^2+^]/[Mn^2+^] ratio for both mutants appears to be about twice as high compared to the WT control (80/1 versus 40/1) indicating alterations of the donor side polypeptide structure have a differential effect with the demand for Ca^2+^ being higher and/or a lower than the demand for Mn^2+^ **(Fig. 4)**. Interestingly, both mutants showed somewhat lower quantum efficiency, *Φ*_*PA*_, compared to WT control, however, at higher [Ca^2+^], the yield continued increasing without saturation or decline through the entire flash sequence, suggesting that while the demand for Ca^2+^ was higher, fewer centers were lost to photoinactivation (compare with WT in Fig. 2). Accordingly, Δ*psbO* and *27OE* appeared less prone to photoinactivation (*Φ*_*PI*_) compared to the WT control **(Fig. 4, S9-S12)** suggesting the open configuration diminishes the tendency for Mn^2+^ to occupy the Ca^2+^ effector site, but with the optimum [Ca^2+^] shifted higher. These observations are consistent with increased diffusion of the metal cofactors inside apo-PSII with proportionally increased exchange rates for Ca^2+^ at its effector site. Both mutants do not show strong correlation between [Ca^2+^] and quantum efficiency of photoactivation **(Fig. 4C)**, which is similar to WT control preparations at the lower, 250µM, [Mn^2+^] tested, but not the higher 500µM, [Mn^2+^]. Photoinactivation rate **(Fig. 4D)** is negligible for both mutant strains. Apparently, increased diffusional access allows Ca^2+^ to more effectively prevent the inhibitory effect of Mn^2+^ on photoassembly. Notably, Δ*psbO* and *27OE* show different Mn dependent photoactivation behavior compared to WT control. While *27OE* has similar to WT control requirement for Mn^2+^ ions, with the optimum at 250µM at 10mM Ca^2+^, Δ*psbO* has lower requirement for available Mn^2+^ and shows maximal yield of photoactivation in the range of concentrations from 50µM to 500µM **(Fig. S9-S12)**.

**Figure 4.**
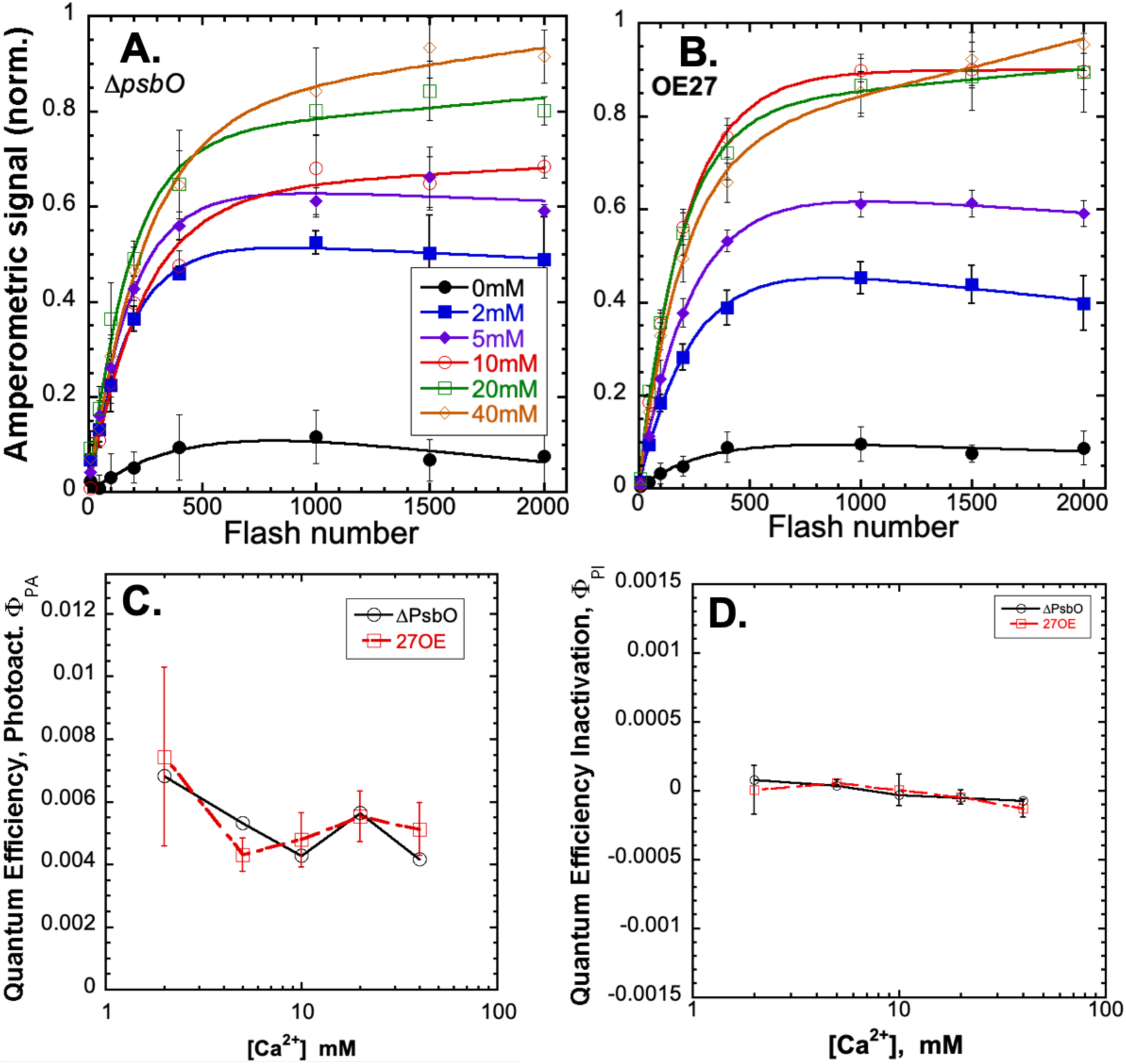
Calcium dependence of photoactivation under train of single turnover flashes of HA-extracted thylakoid membranes from Δ*psbO* (**Panel A**) and *27OE* (**Panel B)** at 0mM (black closed circle), 2mM (blue closed square), 5mM (purple closed diamond), 10mM (red open circle), 20mM (green open square), and 40mM (orange open diamond) of CaCl_2_ combined with 250 μM. **Panel C:** Overall quantum efficiency of photoactivation (*Φ*_*PA*_**) Panel D:** quantum efficiency of inactivation (*Φ*_*PI*_), respectively, in Δ*psbO* (black circle) and 27OE (red square) membranes. Data were fit to equation 1 for parameter estimation (see text for details). Error bars represent standard deviation n ≧3.

As with the WT, the flash interval experiment shows that at longer intervals between flashes, Ca^2+^ stabilizes assembly intermediates **(Fig. 5)** as reflected in the slower decay constant, *k*_D_ **(Table 1)**. However, *27OE* decreases in *k*_D_ overall, further slowing the decay with the increase of [Ca^2+^], suggesting a possible role for Psb27 in stabilizing the photoactivation intermediates **“B”** and/or **“C” (Fig 1)**, perhaps by stabilizing the binding of Ca^2+^ at its effector site. Additionally, the dark rearrangement, *k*_A_, is generally slower than the WT for both Δ*psbO* and *27OE* **(Tables 1)**, although higher [Ca^2+^] does not further slow the dark rearrangement in *27OE* as it does for WT and Δ*psbO*. Apparently both high [Ca^2+^] and the more open configuration of apo-PSII leads to a slower dark rearrangement, *k*_A_ **(Table 1)** yet promotes better exchange of Ca^2+^ and prevents the decay of the photoactivation intermediates **(Tables 1)**. It is worth noting that *27OE* shows higher quantum efficiency, *Φ*_*PA*_, in comparison with Δ*psbO* **(Fig. 4C)**. Taken together, the results suggest that Psb27 enhances the stabilization of intermediates with accessibility of the site of cluster assembly, especially for Ca^2+^ ions, thereby shifting the optimal ratio of Mn^2+^ and Ca^2+^. As discussed, given the known characteristics of the interaction of Psb27 with apo-PSII, (35) this facilitation of photoactivation likely occurs allosterically via its interaction with the luminal E-loop of CP43, which besides D1, provides a ligand to the mature Mn_4_CaO_5_.

**Figure 5.**
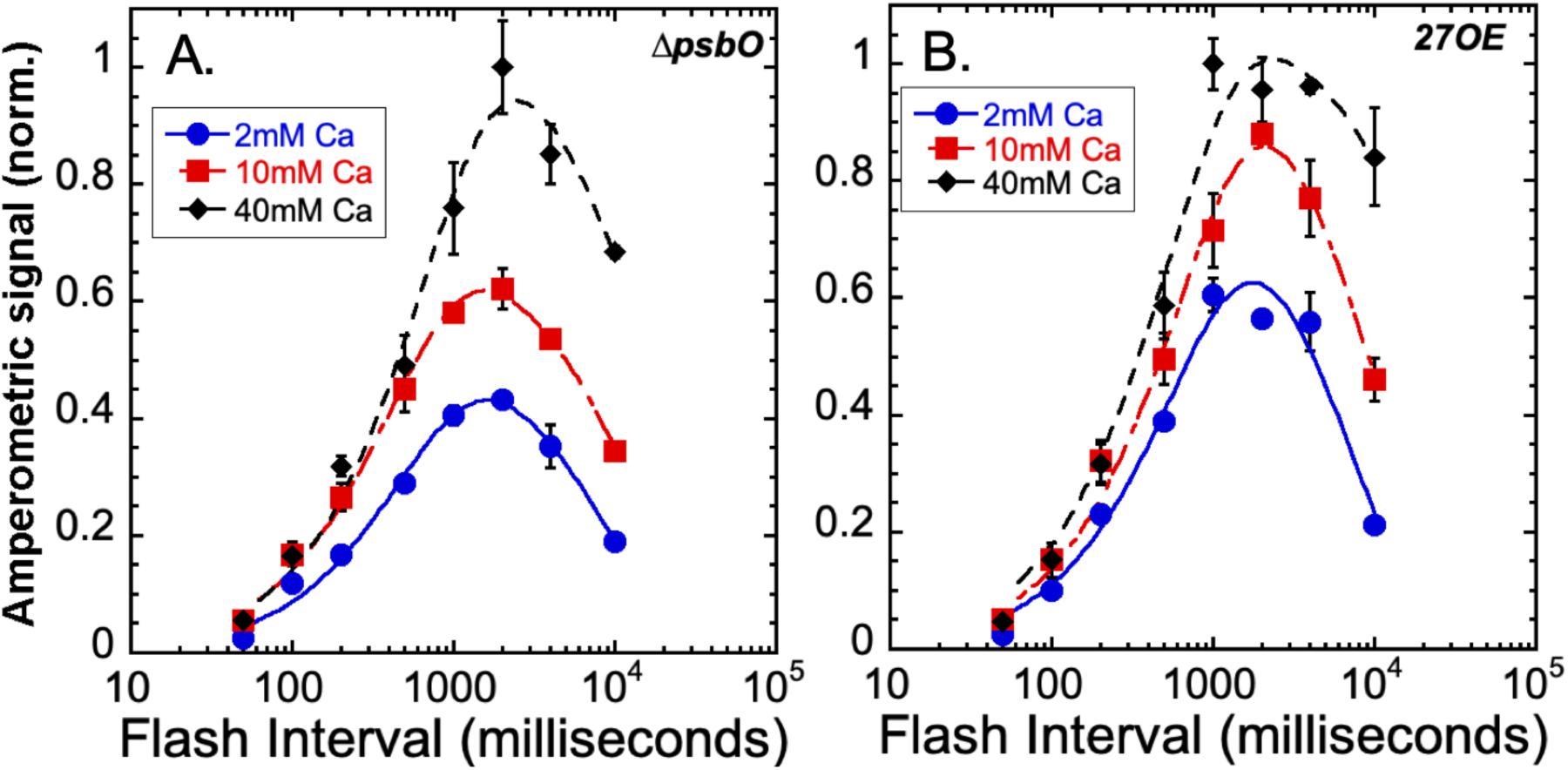
Photoactivation yields as a function of flash interval at different calcium concentrations. Sequences of 150 Xe flashes applied at different flash intervals to HA-extracted thylakoid membranes from Δ*psbO* (**Panel A**) and *27OE* (**Panel B**) with O_2_ yields measured on a bare platinum electrode. Plots correspond to samples containing 2mM (blue circles), 10mM (red squares), and 40mM (black diamond) of CaCl_2_ a fixed [Mn^2+^] = 250 μM. Error bars represent SD with n ≧3. Data was fit to equation 2 (see main text) to estimate the dark rearrangement constant, k_A_, and the decay of photoactivation intermediates “B and C”, k_D_ (see Fig. 1 for kinetic model and **Table 1** for estimated values for k_A_ and k_D_).

**Figure 5.**
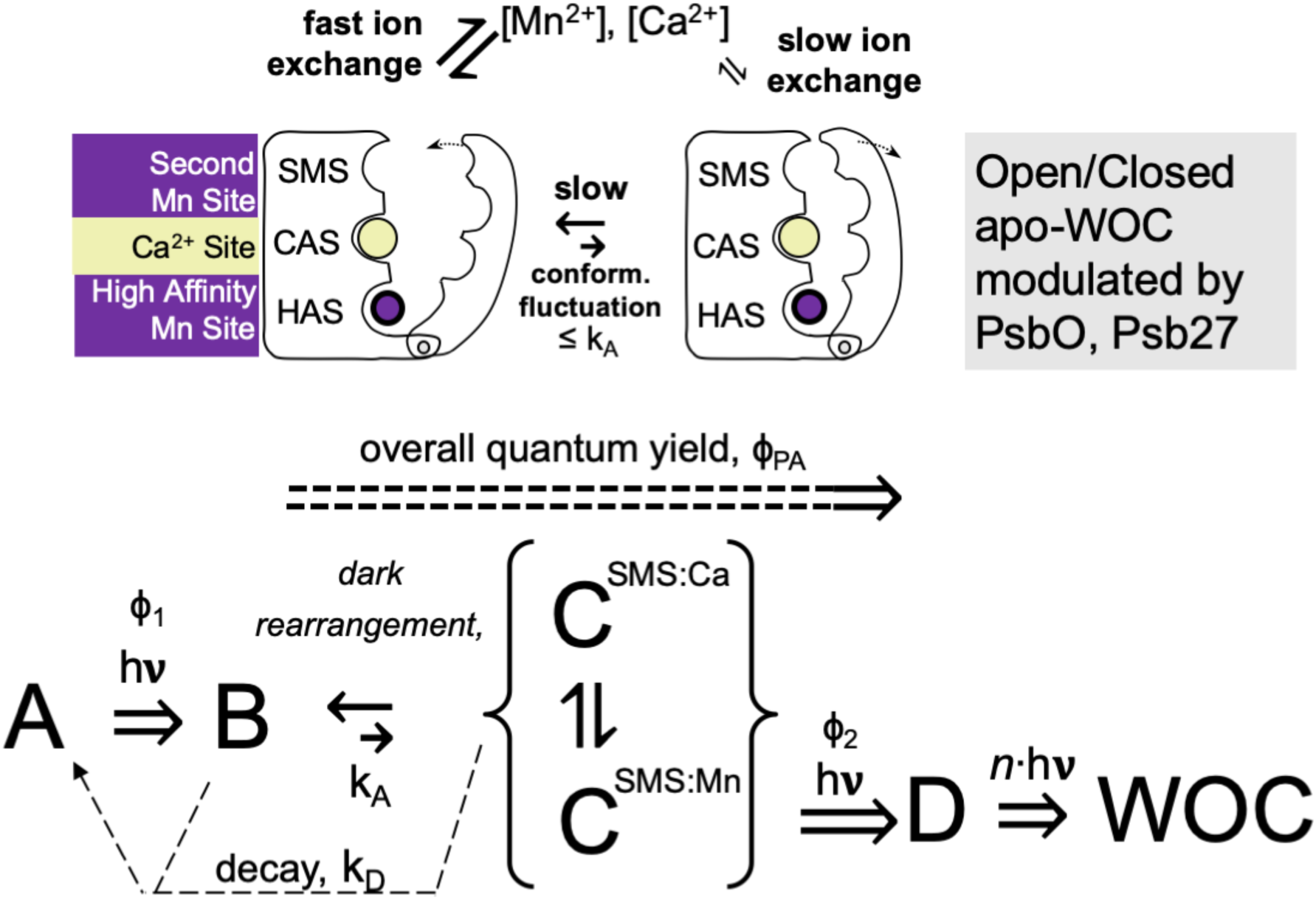
Schematic model of the three main metal binding sites discussed and a minimal model of proposed conformational change and ion exchange accounting for the results.

## Discussion

### How Ca2+ stabilizes photoactivation intermediates yet retards the dark rearrangement

Our results show for the first time that Ca^2+^ stabilizes the assembly intermediates of photoactivation, a feature especially evident at long intervals between photoactivating flashes **(Fig. 3 & 5)**. Moreover, the results also show for the first time that excess Ca^2+^ slows down the already very slow dark rearrangement (**B→C)**, an effect that is enhanced by the more open configuration of the apo-PSII assembly site in the mutants where greater ion exchange occurs **(Fig. 3 & 5, Tables 1)**. According to the two-quantum model of photoactivation (8), a stable intermediate **“D”** is formed by two light-dependent steps separated by a light-independent ‘rearrangement’ (**Fig. 1)**. The first light activated step (**A⇒B**) involves photooxidation of a single Mn^2+^ ion (12, 13) bound as hydroxide [(Mn^2+^)-OH^+1^] (14) at the HAS and occurs with high quantum efficiency (13, 15, 42, 43). The concurrent binding of Ca^2+^ at its adjacent effector site does not affect the affinity of the binding of this Mn^2+^, but upon photooxidation of Mn^2+^ in the presence of bound Ca^2+^, the bridging species, [Mn^3+^-(OH^-^)-Ca^2+^] ⇌[Mn^3+^-(O^2-^)-Ca^2+^] is produced, which is thought to facilitate the next steps (23). This fast, high yield initial photooxidation is followed by a remarkably slow (100-400 ms) rearrangement (**B→C)**, that involves a protein conformational change and/or ion relocation. Only then can the second charge separation become effective (**C⇒D**), photooxidizing a second Mn^2+^, and thereby trapping the first stable intermediate **“D”**. Current models suggest this first stable intermediate, **“D”**, is a binuclear (Mn^3+^)_2_-(di-μ-oxo) bridged structure possibly corresponding to the binuclear di-μ-oxo bridged structure produced by partial disassembly of the intact Mn_4_CaO_5_ using reducing agents (26, 28) or thermal (44) treatment. Based upon the findings that *Φ*_*PA*_ at lower [Mn^2+^] is independent of [Ca^2+^] (**Fig. 1**C) combined with estimates of Mn^2+^ affinity at the HAS (12), we infer that the competition between Ca^2+^ for a critical Mn^2+^ site occurs not at the HAS, but rather at a second Mn site (**SMS**) involved in the photoactivation pathway, and that excess Ca^2+^, while protecting from photoinactivation, inhibits assembly due to occupation of the SMS preventing photooxidation of the second Mn^2+^ (**C⇒D**). The present experiments cannot discriminate what the site is, though it is reasonable to suppose it to involve ligands that form with the other Mn1-Mn3 positions (**Fig. 1C**, orange spheres) and it does not exclude the possibility, for example, that it is a ligation site that the initially photooxidized Mn^2+^ (**A⇒B**) at the HAS relocates to, according to the ‘translocation model’(17, 45).

The stabilization of intermediates (**B, C)** by Ca^2+^ is most simply explained in the same way that Ca^2+^ prevents the photoinactivation of apo-PSII. Photoinactivation is due to incorrect occupation of the Ca^2+^ effector site by Mn^2+^ evident at sub-optimal Ca^2+^/Mn^2+^ ratios, which can even occur in intact PSII (**Fig. S7**) in line with the proposed ‘gate-keeper’ function of Ca^2+^ in the assembled Mn_4_CaO_5_ (26, 28, 39). Furthermore, Ca^2+^ tends to be lost from its effector site due to the electrostatic repulsion occurring upon oxidation of Y_Z_, and the associated charge increase (29, 38). Although the mechanism of Mn-dependent inactivation is not clear, we have concluded that occupation of the Ca^2+^ effector site by Mn^2+^ correlates with photoinactivation as well as ‘misses’ due to the requirement for Ca^2+^ to form the bridging species with Mn at the HAS that is necessary to proceed (23). Therefore, the stabilization of photoactivation intermediates by Ca^2+^ is probably due to this same protective effect and the occupation of the Ca^2+^ effector site by Ca^2+^ and is important throughout the duration of the dark rearrangement (**B→C)** to prevent decay during that process. The observation that both Δ*psbO* and *27OE* each exhibit minimal photoinactivation and enhanced stabilization of intermediates suggests that ion exchange at the Ca^2+^ site is rapid in the more open configuration and this enables centers that have lost the Ca^2+^ ion to rapidly reacquire a replacement.

Besides protecting intermediates, Ca^2+^ also slows the dark rearrangement **(Fig. 3 & 5, Table 1).** At supra-optimal concentrations, Ca^2+^ appears to block the necessary binding of the second Mn^2+^, at the second Mn-binding site, SMS, as already noted. While sufficient concentrations of Ca^2+^ increase the chance of successful formation of intermediate **“C”**, an excess of Ca^2+^ competes with the binding of the second Mn^2+^ binding site, thereby delaying the time before the second Mn^2+^ ion can occupy the SMS for photooxidation, forming the first stable intermediate **“D”**, which is predicted to be a binuclear (Mn^3+^)_2_-(di-μ-oxo) bridged structure. However, if Ca^2+^ competitively occupies the second Mn^2+^ site, then the rearranged configuration (discussed below) cannot be productively converted due to the absence of the second Mn^2+^ needed for the second light step to occur (**C⇒D)**. This competitive inhibition extends the time before the rearranged state can be trapped, at least until the competing Ca^2+^ is replaced with Mn^2+^ enabling the photooxidative formation of stable intermediate **“D”**. This proposed model suggests that the dark rearrangement consists of a molecular reorganization (e.g. conformational change or ion relocation) that is only fruitful if a second Mn^2+^ bound at its correct site.

### How does Psb27 facilitate photoactivation?

It has been shown that Psb27 is the vital player in PSII repair and assembly of the Mn_4_CaO_5_ cluster of PSII that provides greater accessibility to the site of Mn-cluster assembly (31, 32, 35, 46-48). We find that Psb27 facilitates the photoactivation of the WOC in a more complex manner than simply displacing extrinsic polypeptides from apo-PSII. Both Δ*psbO* and *27OE* have increased Mn^2+^ and Ca^2+^ access to the apo-PSII consistent with expectations (reviewed in (25)) and also both mutations produce shifts towards higher optimal [Ca^2+^]/[Mn^2+^] ratios. In the absence of extrinsic proteins, light induces the loss of Ca^2+^ from its binding site in the intact complex and one of the main functions of the extrinsic proteins is to retain the ion (20, 29, 49, 50). Therefore, the shift to higher optimal [Ca^2+^]/[Mn^2+^] ratios is likely because without the retention of Ca^2+^ in the vicinity of the assembly site, relatively higher [Ca^2+^] is required. Therefore, to mitigate the inhibitory effect of high [Mn^2+^], apo-PSII requires more Ca^2+^ ions available to prevent occupation of the Ca^2+^ effector site by Mn^2+^. Both Δ*psbO* and *27OE* mutants could be photoactivated without the concurrent inactivation **(Fig. 4)**, unlike the WT control, which suggests that the increased exchangeability of Ca^2+^ is important for diminishing photoinactivation.

However, if the function of Psb27 only increased exchangeability of Ca^2+^, then Δ*psbO* and *27OE* would have similar phenotypes in regard to photoactivation. This was not the case and *27OE* provides additional support of photoactivation: *27OE* exhibits a remarkable stabilization at high [Ca^2+^], yet does not exhibit a proportionally dramatic increase in the dark rearrangement time at the highest [Ca^2+^] as seen with Δ*psbO* **(Table S1)**. Additionally, it has a greater ability to sustain high yields of photoactivation at moderate [Ca^2+^] compared to Δ*psbO* **(Fig. 4)**. Chemical crosslinking indicates that Psb27 docks to the outer face (distal to the Mn-assembly site) of the E-loop of CP43 and exerts its effects, including the weakened binding of extrinsic proteins allosterically (35). This suggests that the Psb27-E-loop interaction stabilizes a structural arrangement that: 1.) enhances the selectivity of the second Mn^2+^ photooxidation site versus the competitive binding of Ca^2+^ at the second Mn site 2.) stabilizes the binding of Ca^2+^ to the Ca-effector site, and 3.) maintains an open configuration that enables rapid rebinding of Ca^2+^ if the ion is lost from the Ca-effector site and/or facilitates the exchange of metals in malformed metal centers.

### What is the dark rearrangement?

The utilization of single turnover flashes during the flash interval experiment ensures that the dark rearrangement (**B→C)** estimates the rate, k_A_, of a molecular process proposed to be a conformational change (22, 37, 40, 51) or the relocation of a bound ion (17, 45). Once the rearrangement has occurred, the labile configuration, can be trapped by a second photooxidation of the second Mn^2+^, to produce the first stable intermediate, “**D**”. Based upon the observation that Psb27 allosterically modulates the assembly of the Mn_4_CaO_5_, likely through its known interaction with the CP43-E-loop (35, 36), we suppose that the mobility of the E-loop is involved in the Mn_4_CaO_5_ assembly process, possibly as part of the dark rearrangement. The E-loop is a globular domain situated between transmembrane helices 5 and 6 of CP43 and directly interacts with the assembled Mn_4_CaO_5_, including a bridging carboxylate ligand to Mn2 and Mn3 via CP43-Glu354 (5). Moreover, it directly contacts the C-terminal domain of the D1-protein, which contains amino acids coordinating the Mn_4_CaO_5_, and notably provides a ligand to the Ca^2+^ and Mn1 via the carboxyl group of its C-terminus. Thus, movements of the E-loop are likely coupled to movements of the C-terminal domain, and *vice versa*. The E-loop is connected to the membrane intrinsic portion of the protein via a narrow neck that potentially gives the domain mobility to accommodate the assembly/disassembly of the WOC by rocking in and out of contact with the Mn assembly site (18) and these postulated structural fluctuations may be part of the dark rearrangement **(Fig. 6)**. The increased accessibility in Δ*psbO* cannot be explained by steric covering of the apo-WOC and instead the open configuration must relate to the fact that PsbO forms a structural bridge between the E-loop and parts of the D1 protein including the C-terminal domain (**Fig. 1B**). Thus, the more open configuration due to the loss of PsbO is likely due to enhanced mobility of the E-loop and associated structural elements enabling the open configuration. Recent cryoEM structure data has provided the first direct evidence for the alternate position of the E-loop (*Gisriel, Brudvig pers. comm*). A more open position of the E-loop may be stabilized by Psb27, so that it still may fluctuate, but in a range that optimizes assembly, in contrast to the loss of PsbO, where the more open configuration is evident, but not the other kinetic features, which are absent. However, it seems unlikely that the predicted mobility of the apo-WOC would alone explain the conformational rearrangement because the rate, k_A_, slows down rather than speeds up in the mutants and it is strongly affected by [Ca^2+^]. So, while conformational mobility of the E-loop and the associated D1-carboxy terminus is a reasonable explanation for the more open structure and thereby minimizes photoinactivation as discussed above, how conformational mobility would also slow the rearrangement process is not readily evident. One possibility is that the increased mobility also involves a change in the distribution of conformational states towards the more open configuration away from the closed configuration. Thus, if the dark rearrangement corresponds to conformational fluctuations alternating between the open and closed configurations, where the open configuration is energetically favored in Δ*psbO* and *27OE*, then the frequency with which the closed configuration is formed would occur less frequently. This combined with the need for simultaneous binding of the second Mn^2+^ (and not competitive Ca^2+^) to trap **“D”**, could explain the how supra-optimal [Ca^2+^] also slows the rearrangement. In this model, the rearrangement rate reflects the frequency of occurrence that the first photoinitiated intermediate, **“B”** containing Mn^3+^ has conformationally rearranged and the SMS site is actually occupied by a Mn^2+^ ion **(Fig. 6)**. At high [Ca^2+^]/[Mn^2+^] ratios, the conformational rearrangement may occur, but the correct occupancy of the SMS may not have occurred due to competitive displacement and the additional time for the extended re-arrangement period reflects the time it takes for the competing Ca^2+^ to exchange with the necessary Mn^2+^.

## Materials and Methods

### Strains and Growth conditions

The glucose-tolerant *Synechocystis* sp. PCC6803 control strain (WT-control) expressing only the wild-type *psbA2* gene and having a hexa-histidine tag fused to the carboxyl-terminus of CP47 was maintained in BG-11 medium as described previously (52). Experimental cultures were grown in flat 1L tissue culture flasks in 800 mL BG-11 media buffered with 10 mM HEPES-NaOH pH 8.0 (HBG-11) supplemented with 5mM glucose (Sigma) under a PFD (photon flux density) of ∼80 µmol m^-2^ s^-1^ at 30 °C. Cultures were bubbled with filter sterilized air. Light intensity measurements were made with a Waltz light meter (Germany).

### Mutant Strain Generation

The Δ*psbO* deletion mutant was constructed previously and involved replacement of the *psbO* coding sequence with a Sp^r^/Sm^r^ antibiotic resistance gene (41). For overexpression of Psb27 gene in *Synechocystis*, the chromosomal locus comprising the open reading frame slr1645 (Psb27) was amplified by PCR **(Table S2)** using Herculase II Phusion DNA polymerase (Agilent USA) and integrated at a neutral site within the *Synechocystis* genome between ORFs slr1169 and slr1285 (**Fig. S1**). Transformation was carried as described previously using selective agar plates containing 5mM glucose, 10µM DCMU, and 12.5µg/mL spectinomycin followed by further selection at 25µg/mL spectinomycin (53).

### Preparation of Mn-depleted thylakoid membranes

Cultures (2.4 L) were harvested in early stationary phase (OD_750nm_ ∼1.8-2.2) and exhibited variable Chl fluorescence ((*F*_m_ - *F*_0_)/*F*_0_) values >0.5, as measured with a Photon Systems Instruments (PSI) Fluorometer FL 3500. Cells were collected by centrifugation at 25°C at 6,000*g* (Sorvall, F-9 rotor) for 15 min and gently resuspended with a minimal volume of H20BG-11(same as growth media, but with 20mM buffer) using a paintbrush. The cell suspension volume was expanded to ∼200 mL with additional HBG-11. The washed cells were centrifuged again at 25 °C at 10200*g* for 5 min (Sorvall, F-14 rotor). The pelleted cells were resuspended as before in HBG-11 medium, and the suspension was adjusted to a Chl concentration of 100 μg mL^-1^ Chl. Hydroxylamine (HA) was added to 1 mM from a freshly prepared 400 mM stock, and the treated suspension was incubated for 12 min in the darkness with rotary agitation (200 rpm) at room temperature. Maintaining complete darkness, the HA-extracted cells were then washed by resuspension with 100 ml HBG-11 and rotary agitation (200 rpm) for 5 min before pelleting again. This washing step was repeated four more times with the aim of depleting residual HA. Finally, the cells were resuspended in 120 ml of breaking buffer (1.2M betaine, 50mM MES-NaOH (pH 6.0), 10%(v/v) glycerol, and 5mM MgCl_2_) that was prepared with ultrapure reagents and Chelex-100 (Bio-Rad)-treated solutions and incubated in the dark on ice for 1 hour. Cells were pelleted at 10,200*g* for 5 min and resuspended to a total volume of 14 mL. Prior cell breakage, 1 mM benzamidine, 1 mM ε-amino-n-caproic acid, 1 mM PMSF, and 0.05 mg/mL DNase I were added. Cells were broken by four cycles of 5sec ON and 5min OFF in an ice-cooled glass bead homogenizer (Bead-Beater, BioSpec Products, Bartlesville, OK). After breakage the sample was centrifuged at 3,600g for 10 min to pellet unbroken cells and cell debris. Thylakoids were obtained from the supernatant cell homogenate by ultracentrifugation (35 min at 40,000 rpm in a Beckman Ti45 rotor) and thylakoid-containing pellets were suspended in Chelex-100-treated breaking buffer to a concentration of 1.0-1.5 mg of Chl mL^-1^ with [Chl] determined from methanol extracts according to (54). Concentrated thylakoid membranes were flash frozen as 100 µL aliquots in liquid nitrogen and stored at −80°C.

### Photoactivation of HA-extracted membranes

HA-extracted membranes were photoactivated either in suspension for subsequent assay for restoration of O_2_ evolution detected using a Clark-type electrode or directly on a bare platinum electrode that permits the centrifugal deposition of samples upon the electrode surface (15) (**Fig. S2**). Flash illumination under each mode of photoactivation was provided using an EG&G xenon flash lamp. The buffer was supplemented with various concentrations of cations CaCl_2_, MnCl_2_, SrCl_2_, and MgCl_2_ to examine their role in photoactivation for the role of cations on photoactivation. At the completion of the photoactivation flash treatment on the bare platinum electrode, flash O_2_ yields were recorded as described previously (55, 56). Photoactivation was also monitored based upon light-saturated steady-state rates of O_2_ evolution of the photoactivated thylakoid membranes, using a Clark-type electrode (9, 12, 40). HA-extracted thylakoid samples containing 40 µg of Chl were suspended in 400 µL of in Chelex-100-treated photoactivation buffer in a modified aluminum weight cup with a stirring bar. A series of single-turnover xenon flashes was given to the stirring thylakoid suspension and O_2_ evolution rates were in O_2_ evolution buffer (50 mM MES-NaOH, 25 mM CaCl_2_, 10 mM NaCl, and 1M sucrose pH 6.5) at a light intensity of 2500 µEm^-2^ s^-1^ in the presence of of 1mM 2,6 dichloro-*p*-benzoquinone (DCBQ) and 2.5mM potassium ferricyanide K_3_[Fe(CN)_6_].

### SDS-PAGE and Immunoblot analysis

Membrane samples obtained as above were solubilized by addition of 2% SDS, 5mM dithiothreitol, heated at 65 °C for 10 min, and separated on SDS-PAGE (12%). Protein content was transferred to a PVDF membrane using a Bio-Rad semi-dry apparatus, and membrane was stained with 0.5 % Ponceau S (Sigma) to verify equal loading. 5% BSA solution was used as a blocking agent. Blots were probed against Psb27 (1:750 dilution, a kind gift from Julian Eaton-Rye) with anti-rabbit HRP-conjugated goat antibody (1:3000, Bio-Rad) as a secondary antibody. Color Development was obtained using the chromogenic substrate 4-chloro-1-naphthol (4CN, Bio-Rad) and H_2_O_2_.

## Acknowledgments

The authors wish to thank Prof. Charles Yocum for stimulating discussions and great technical advice. We thank Rachel Martin for contributions to the early phases of this study. This work was funded by the National Science Foundation NSF MCB-1716408.

