## Supplemental Information for "The role of Ca^2+^ and protein scaffolding in the formation in nature’s water oxidizing complex"

*Short Title: Assembly of water oxidation catalyst*

Anton P. Avramov, Hong J. Hwang, and Robert L. Burnap

Department of Microbiology & Molecular Genetics, Oklahoma State University, Stillwater, USA.

Corresponding Author: Robert L. Burnap, Department of Microbiology and Molecular Genetics 307 Life Sciences East, Oklahoma State University, Stillwater, OK 74078. Telephone: 405-744-7445, Fax: 405-744-6790,

**Table S1**. Photoactivation of hydroxylamine (HA)-extracted and non-extracted membrane samples obtained from *Synechocystis* WT control*,* ∆*psbO* and *27OE* strains. Photoactivation was produced by application of 1000 saturating single turnover xenon flashes in HA-extracted thylakoid samples containing 40 µg of Chl, which were suspended in 400 µL of in Chelex-100-treated photoactivation buffer in a modified aluminum weight cup with a stirring bar. Single-turnover xenon flashes were given to the stirring thylakoid suspension. Maximal rate of oxygen evolution were determined for both HA-extracted and unextracted samples containing 10 µg of chlorophyll were used to measure oxygen evolution rate at 30 °C using Clark-type electrode in a buffer containing 50 mM MES-NaOH, 25 mM $CaCl_2$, 10 mM NaCl, 1M sucrose pH 6.5, 1mM 2,6 dichloro-*p*-benzoquinone (DCBQ) and 2.5mM potassium ferricyanide $K_3[Fe\ (CN)_6]$.

| Strain | Non-extracted thylakoid membranes, µmol $O_2$ (mg Chl)$^{-1}$ h$^{-1}$ | Apo-PSII µmol $O_2$ (mg Chl)$^{-1}$ h$^{-1}$ | Recovered rate µmol $O_2$ (mg Chl)$^{-1}$ h$^{-1}$ | % recovery |
|---|---|---|---|---|
| ***WT*** | 575.6±13.5 | 12.1±7 | 224±5.3 | 38.9 % |
| ***27OE*** | 330±24.8 | 17± 8.4 | 193.6±8.5 | 58.4 % |
| **∆*psbO*** | 254.6±17.3 | 5± 3 | 146.7±20.4 | 57.6 % |

**Table S2:** DNA Primers used for the cloning and overexpression of the psb27 gene.

| Primer name | Sequence |
|---|---|
| Psb27_GA_F | TACATAAGGAATTATAACCAATGTCCTTTTTGAAAAATCAGTTGTCACGGC |
| Psb27_GA_R | TTATCAGACCGCTTCTGCGCTCACACGCCCCGTTCAATG |
| slr1169_seq_F | CGATGATGGCGATCGCCAAAAC |
| Sp_cassette_seq _R | GGTGGTAACGGCGCAGTG |
| slr_1285_seq_R | CACCCCCACGCCATCAA |
| Psb27_seq_F | CCACCGGCATCACCTTTGC |

**Figure S1**

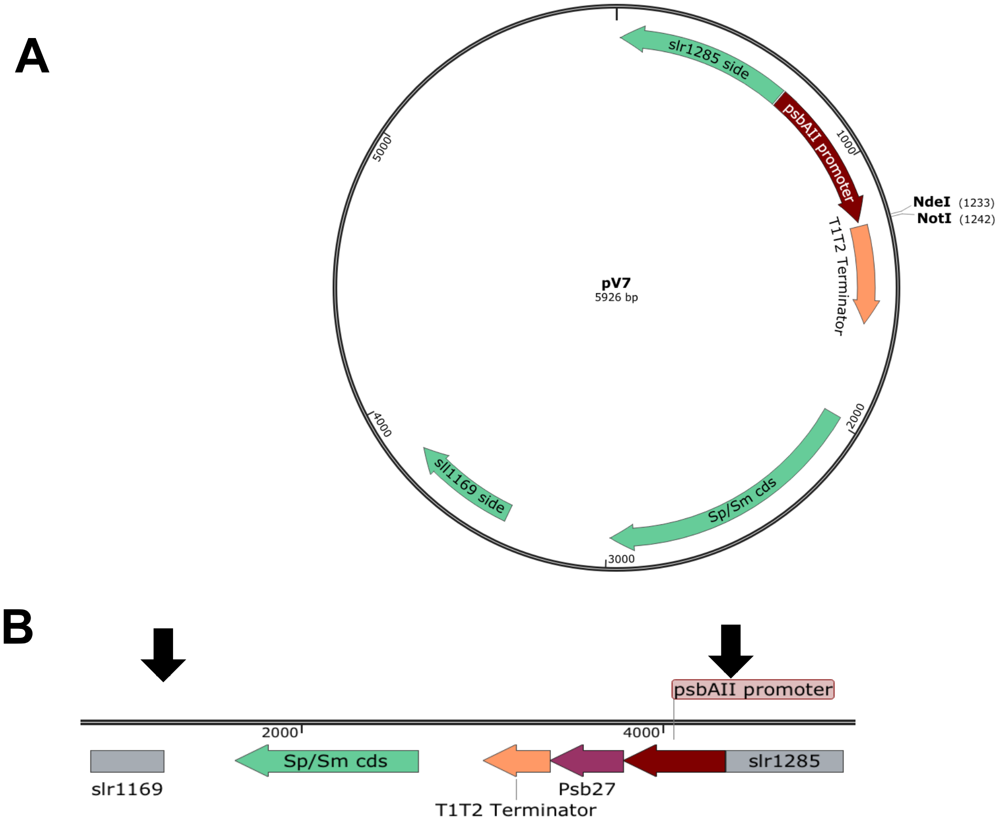


**Figure S1.** The construction of ectopic expression mutant *27OE*. (A) pV7 plasmid used for the mutant construction, (B) integration of Psb27 gene to the *Synechocystis sp. PCC6803* genome. Arrows indicate the insertion of Psb27 gene with psbAII promoter and T1T2 terminator in conjunction with Spectinomycin/Streptomycin antibiotic resistance cassette. The chromosomal locus comprising the open reading frame slr1645 (Psb27) was amplified by PCR **(Table S2)** using Herculase II Phusion DNA polymerase (Agilent USA). The amplified DNA fragment was assembled into pV7 plasmid (a gift from Hong Wang and Wim Vermaas, Arizona State University) between NotI and NdeI restriction sites using the Gibson Assembly technique (38). The pV7 vector is a suicide plasmid that contains DNA sequences that mediate homologous recombination within the *Synechocystis* chromosome and is designed for the introduction of ectopic overexpression cassettes, in this case, the *psb27* gene was placed under the control of the strong *psbAII* promoter. Transformation results in the integration of the expression cassette at a neutral site within the *Synechocystis* genome between ORFs slr1169 and slr1285.

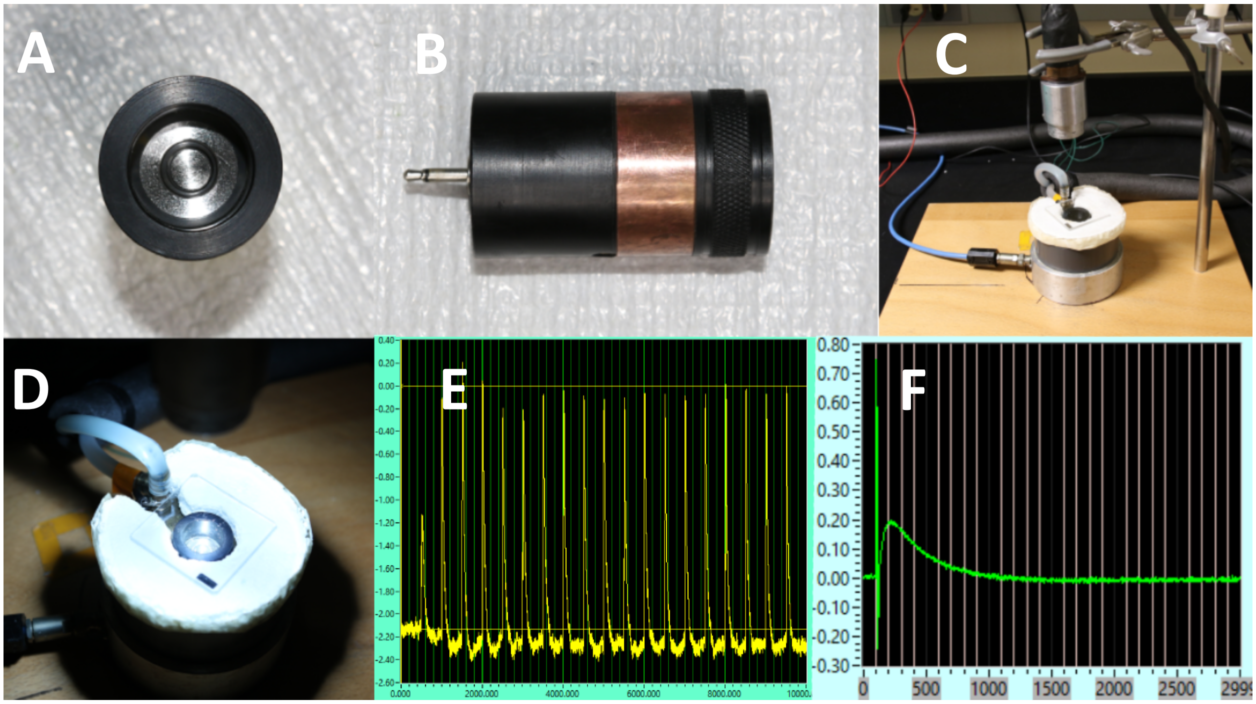


**Figure. S2.** Experimental setup. **A)** and **B)** Bare platinum electrode that permits the centrifugal deposition of the sample upon the electrode surface at 8000g in Thermo Scientific TH-13 rotor. **C)** and **D)** bare platinum electrode with the EG&G xenon flash lamp receiving 4.5 J discharge from 5μF capacitor connected to an EG&G PS-302 triggered power supply with a flash duration of 5μs (full-width, at half-maxima peak intensity), light from the flash lamp was focused vertically down on an electrode. For measurements using a bare platinum electrode, membrane samples containing 3 *μ*g of Chl in 600 *μ*L of a Chelex-100-treated photoactivation buffer consisting of 50 mM MES-NaOH, 5 mM $MgCl_2$, 250 mM NaCl, and 800 mM sucrose (pH 6.5) were centrifugally deposited at 8,000*g* at 25°C for 8 min onto the platinum surface of the electrode in a Sorvall TH-13 swingout rotor. Electrode surface was cleaned prior every measurement with $NaHCO_3$ powder and water. **E)** LabView programming environment (National Instruments, Austin, TX) connected to a NI USB-6361 data acquisition card with internal clocks allowing to collect 10000 measurements per second. **F)** LabView programming environment application for analysis of the complex flash patterns.

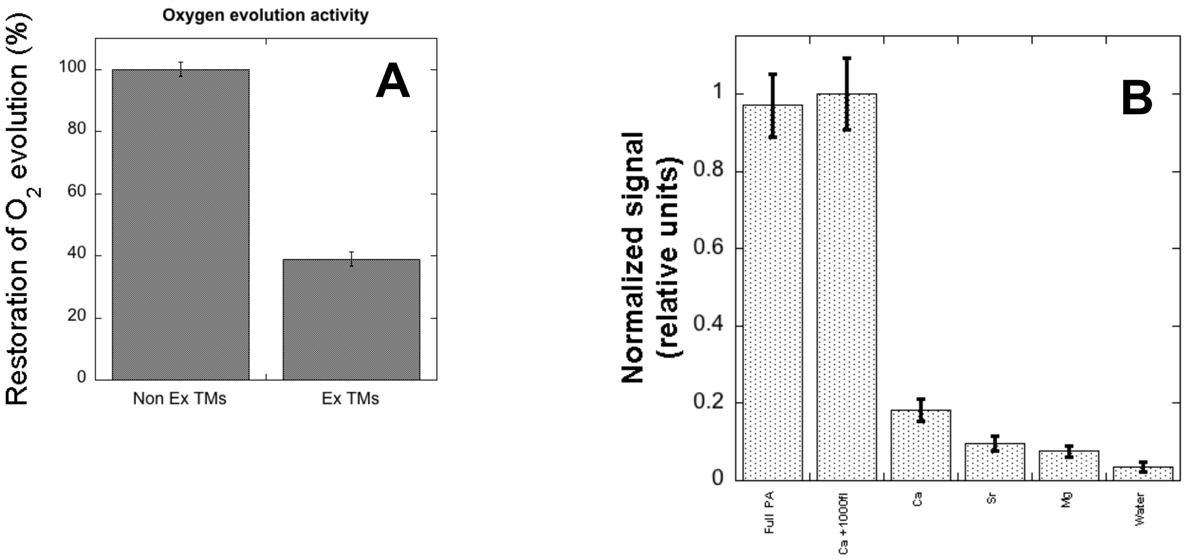


**Figure S3.** Photoactivation of hydroxylamine-extracted thylakoid membranes from WT control (A) Recovery of the light-saturated $O_2$-evolving activity of hydroxylamine-extracted thylakoid membranes exposed to 15 minutes of continuous illumination. (B) Effect of post-adding of divalent cations ***after*** illuminating the sample with 1000 photoactivating Xenon flashes in the presence of 250µM $MnCl_2$. The thylakoid membranes were photoactivated using 1000 single-turnover flashes illumination. The 1st photoactivation was carried out in the presence of $Mn^{2+}$ and then cations were added to the photoactivated sample. After incubating for 10 min in the dark on the ice, the photoactivation activity was monitored by measuring $O_2$ evolution using centrifugal bare platinum electrode using 20 measuring Xenon flashes.

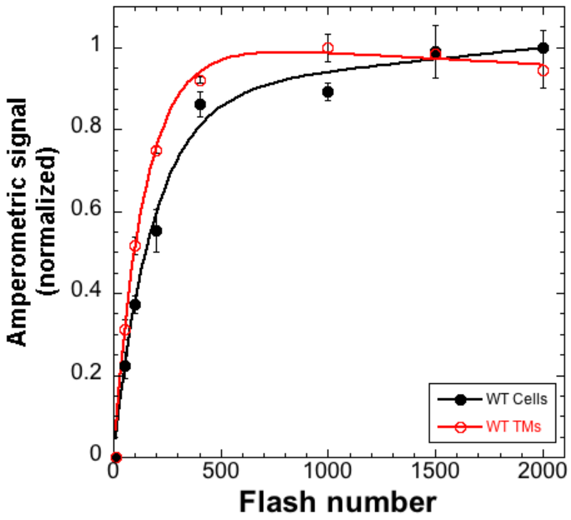


**Figure S4.** Photoactivation kinetics of hydroxylamine-extracted WT control cells (black closed circles) and hydroxylamine-extracted WT control thylakoid membranes (red open circles) as a function of the flash number. Photoactivating flashes were given at uniform interval of 0.5 sec in the presence of 10mM of $CaCl_2$ and 250µM $MnCl_2$. The photoactivation kinetics of cells and thylakoids were measured on Clark-type electrode and Joliot-type electrode, respectively. Obtained data was fit to equation (1, main text). Error bars represent standard deviation $n \geqq 3$.

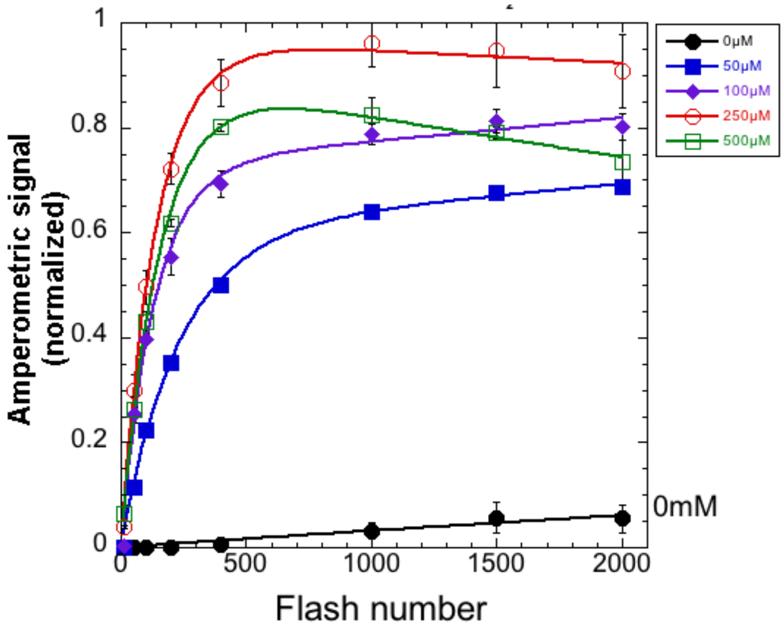


**Figure. S5** Photoactivation of HA-extracted thylakoid membranes from WT control as a function of flash number at different $Mn^{2+}$ concentrations: 0μM (black closed circle), 50μM (blue closed square), 100μM (purple closed rhombus), 250μM (red open circle), and 500μM (green open square) of $MnCl_2$ as a function of the flash number. Development of photoactivated PSII was monitored polarographically in response to photoactivating flashes given with a uniform interval of 0.5 sec at a fixed $[Ca^{2+}]$ = 10 mM. Data were fit to equation (1) Error bars represent standard deviation n ≧ 3.

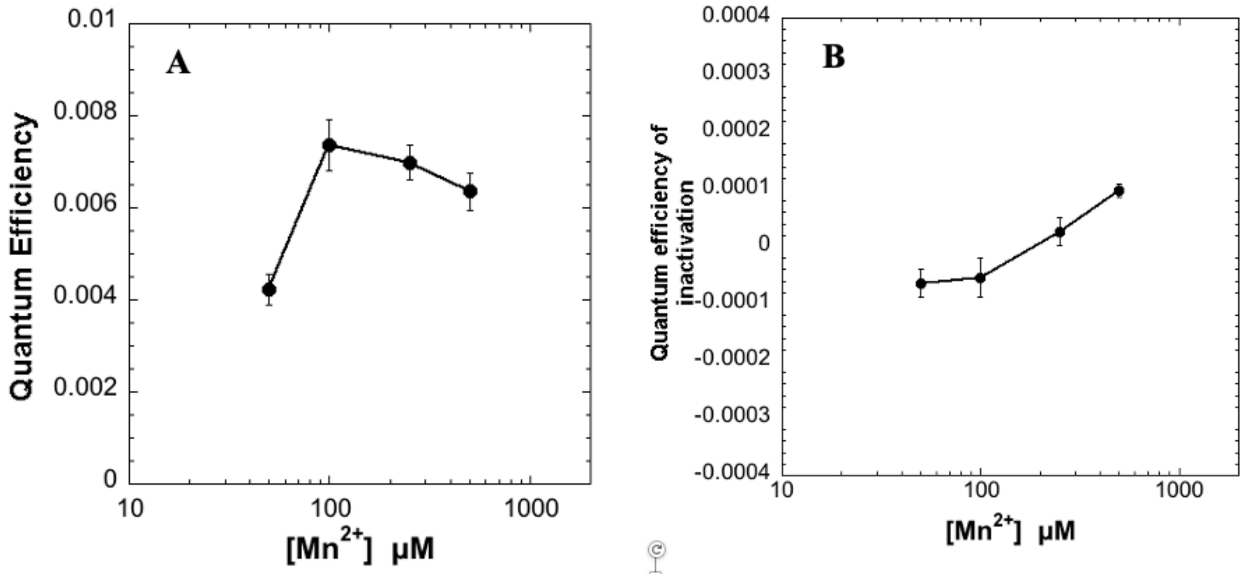


**Figure S6**. Effect of the different concentrations of $MnCl_2$ at a fixed $[Ca^{2+}]$ = 10 mM on (A) Quantum Efficiency and (B) Quantum efficiency of inactivation in WT control. Kinetic values obtained from fits of data in Figure S3 to equation (1). Error bars represent standard deviation n ≧ 3.

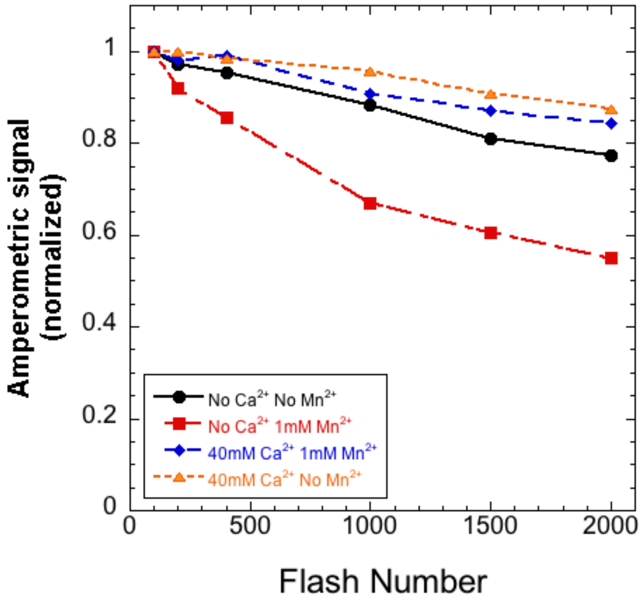


**Fig S7.** $Mn^{2+}$ induced photoinactivation of oxygen evolution in thylakoid membranes from WT control at different ion availability conditions as a function of the flash number with different amounts of $Ca^{2+}$ and $Mn^{2+}$ added to photoactivation buffer. No $Ca^{2+}$ and no $Mn^{2+}$ (black circles), no $Ca^{2+}$ and 1mM $Mn^{2+}$ (red squares), 40mM Ca2+ and 1mM $Mn^{2+}$ (blue rhombus), and 40mM $Ca^{2+}$ and no $Mn^{2+}$ (orange triangles).

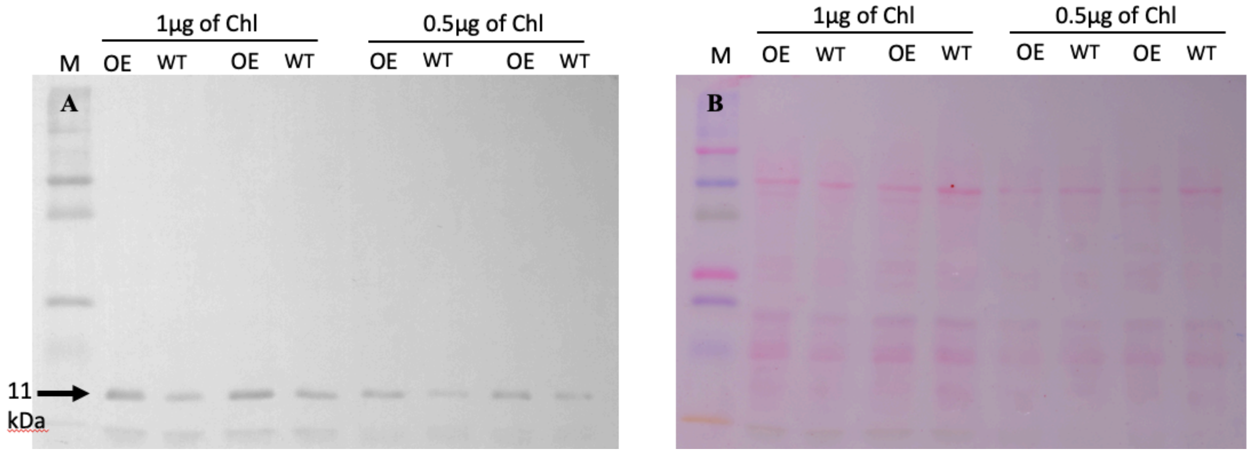


**Figure S8.** A) Immunological detection of Psb27 expression in WT control and *27OE.* Whole cell extracts samples containing 0.5μg or 1 μg Chl were loaded on a 12% SDS-PAGE gel and transferred to a PVDF membrane. B) Previous to detection, the membrane was stained with 0.5% Ponceau S to verify equal loading. Expression of Psb27 (~11kDa) was detected using antibody (a kind gift from Julian Eaton-Rye). A marker was added for molecular weight comparisons (lane M) (Precision Plus Protein Kaleidescope, Bio-Rad).

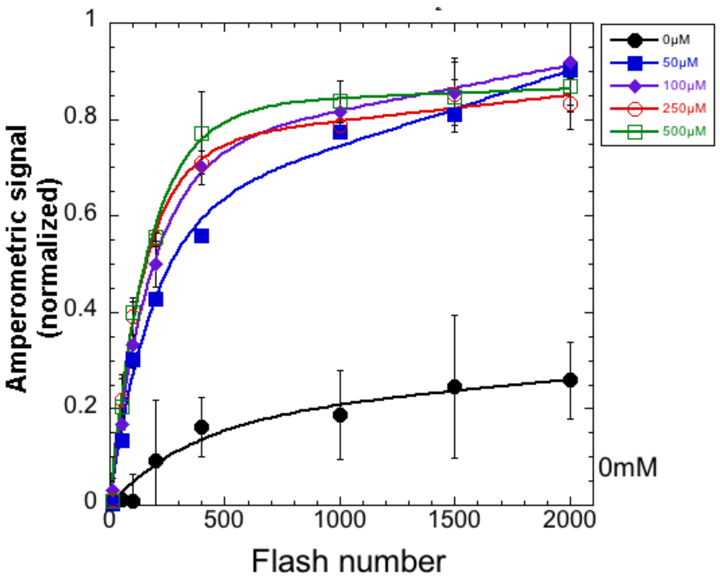


**Fig S9.** Photoactivation of hydroxylamine-extracted thylakoid membranes of Δ*psbO* as a function of flash number at different $Mn^{2+}$ concentrations: 0μM (black closed circle), 50μM (blue closed square), 100μM (purple closed rhombus), 250μM (red open circle), and 500μM (green open square) of $MnCl_2$ as a function of the flash number. Development of photoactivated PSII was monitored polarographically as oxygen yields on a bare platinum electrode in response to photoactivating flashes given with a uniform interval of 0.5 sec at a fixed $[Ca^{2+}]$ = 10 mM. Data were fit to equation (1) describing a monoexponential consumption of apo-PSII and a parallel inactivation process (see text for details). Error bars represent standard deviation n ≧ 3.

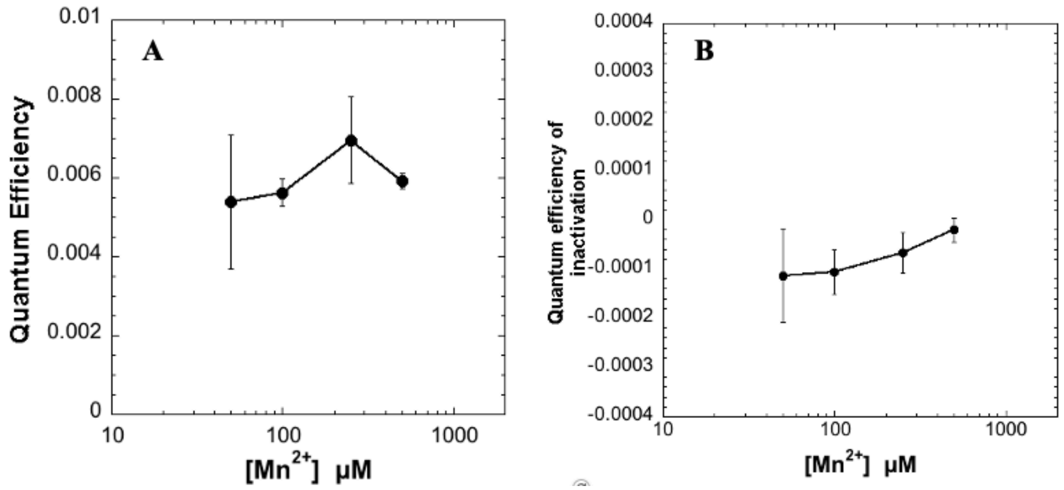


**Figure S10**. Effect of the different concentrations $MnCl_2$ at a fixed $[Ca^{2+}]$ = 10 mM on (A) Quantum Efficiency and (B) Quantum efficiency of inactivation in Δ*psbO*. Kinetic values obtained from fits of data in Figure S9 to equation (1). Error bars represent standard deviation n ≧ 3.

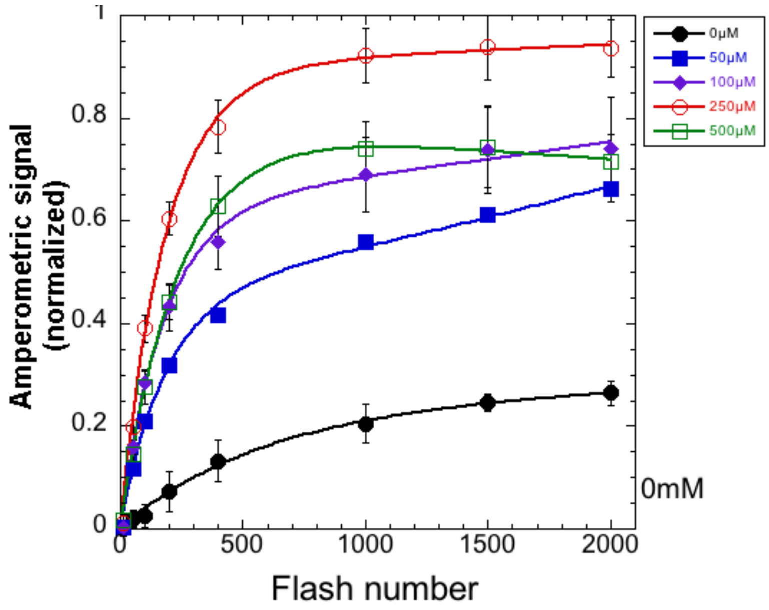


**Fig S11.** Photoactivation of hydroxylamine-extracted thylakoid membranes from *27OE* as a function of flash number at different $Mn^{2+}$ concentrations: 0μM (black closed circle), 50μM (blue closed square), 100μM (purple closed rhombus), 250μM (red open circle), and 500μM (green open square) . Development of photoactivated PSII was monitored polarographically as oxygen yields on a bare platinum electrode in response to photoactivating flashes given with a uniform interval of 0.5 sec at a fixed $[Ca^{2+}]$ = 10 mM. Data were fit to equation (1) (see text for details). Error bars represent standard deviation n ≧ 3.

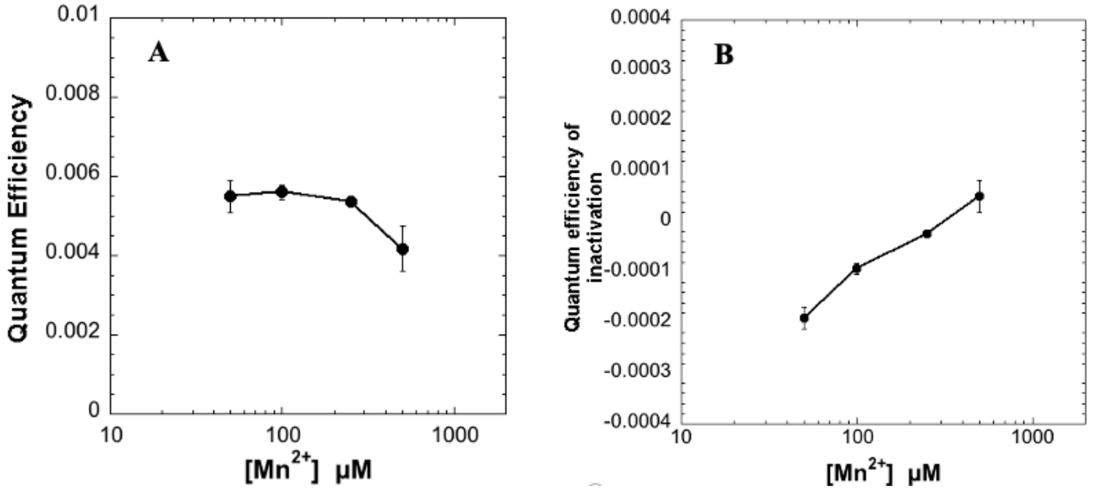


**Fig S12**. Effect of the different concentrations $MnCl_2$ at a fixed $[Ca^{2+}]$ = 10 mM on (A) Quantum Efficiency and (B) Quantum efficiency of inactivation in WT control *27OE*. Kinetic values obtained from fits of data in Figure S9 to equation (1). Error bars represent standard deviation n ≧ 3.
